# The *Plasmodium falciparum* CCCH zinc finger protein MD3 regulates male gametocytogenesis through its interaction with RNA-binding proteins

**DOI:** 10.1101/2023.07.19.549485

**Authors:** Afia Farrukh, Jean Pierre Musabyimana, Ute Distler, Vanessa Jil Mahlich, Julius Mueller, Fabian Bick, Stefan Tenzer, Gabriele Pradel, Che Julius Ngwa

**Affiliations:** Division of Cellular and Applied Infection Biology, Institute of Zoology, RWTH Aachen University, Aachen, Germany; Core Facility for Mass Spectrometry, Institute of Immunology, University Medical Centre of the Johannes Gutenberg University, Mainz, Germany

**Keywords:** *Plasmodium falciparum*, malaria, RNA-binding protein, CCCH zinc finger protein, male gametocytogenesis, exflagellation

## Abstract

Malaria transmission to mosquitoes is dependent on the formation of gametocytes. When fully matured, gametocytes are able to transform into gametes in the mosquito’s midgut, a process accompanied with their egress from the enveloping erythrocyte. Gametocyte maturation and gametogenesis require a well-coordinated gene expression programme that involves a wide spectrum of regulatory proteins, ranging from histone modifiers to transcription factors to RNA-binding proteins. Here, we investigated the role of the CCCH-zinc finger protein MD3 in *P. falciparum* gametocytogenesis. MD3 was originally identified by us as an epigenetically regulated protein of immature gametocytes and recently shown to be involved in male development in a barcode-based screen in *P. berghei*. We here show that MD3 is mainly present in the cytoplasm of immature male *P. falciparum* gametocytes. Parasites deficient of MD3 are impaired in gametocyte maturation and male gametocyte exflagellation. BioID analysis in combination with co-immunoprecipitation assays unveiled an interaction network of MD3 with RNA-binding proteins like PABP1 and ALBA3, with translational initiators, regulators and repressors like elF4G, PUF1, NOT1 and CITH, and with other regulators of gametocytogenesis, including ZNF4, MD1 and GD1. We conclude that MD3 is part of a regulator complex crucial for post-transcriptional fine-tuning of male gametocytogenesis.

## Introduction

Malaria is a life-threatening tropical disease caused by parasites of the genus *Plasmodium*, which accounts for over 247 million infections and 619,000 deaths in 2021 (World malaria report 2022). Among the five species of *Plasmodium* that infect humans, *Plasmodium falciparum* is the most lethal and responsible for the majority of malaria-related deaths particularly in WHO global Africa. To date, malaria eradication is being hampered by the rapid spread of resistance against frontline drugs, limited efficacy of the available malaria vaccine RTS,S and risk of disease aggravation due to climate change (Mora *et al*., 2022). While the clinical symptoms and pathology of malaria are primarily associated with the asexual blood stages of the parasite, the sexual stages, particularly the gametocytes, play a critical role in the transmission of the disease.

*Plasmodium falciparum* gametocytes are sexual precursors of the parasite that develop within the human host’s bloodstream. These highly specialized cells are responsible for the production of male and female gametes following their uptake by the *Anopheles* mosquito vector, a process that requires their egress from the enveloping red blood cell (RBC). Formation and maturation of *P. falciparum* gametocytes in the human blood takes roughly 10 days. Gametocyte development is a tightly regulated process that involves complex molecular mechanisms of gene regulation, which drive the morphological and physiological changes in the gametocytes including sex determination, and which also prime the gametocytes for gametogenesis (reviewed in e.g., Bennink *et al*., 2016). The regulation of gene expression in gametocytes is particularly important to ensure on the one hand sexual commitment, i.e. their differentiation from asexual blood stages and the acquisition of sexual competence, and on the other hand to prepare the parasite for a changing environment during human-to-mosquito transmission.

Sexual commitment occurs in a small fraction of asexual blood-stage parasites and is triggered by environmental cues, such as changes in temperature and nutrient availability. Sexual commitment is accompanied by sex determination of gametocytes and to date, it is unknown how, in the absence of sex chromosomes, two diverging gene expression programs leading to male and female gametocytes are started in the haploid *Plasmodium* parasites. It is meanwhile known that transcriptional control plays a major role in both sexual commitment and sex determination. The gametocyte development protein 1 (GDV1) acts as a key regulator of sexual commitment by facilitating dissociation of heterochromatin protein 1 (HP1) from heterochromatic DNA at the genomic locus encoding AP2-G, the master transcription factor of gametocytogenesis (e.g. Brancucci *et al*., 2014; Coleman *et al*., 2014; Filarsky *et al*., 2018; Usui *et al*., 2019). AP2-G belongs to a group of ApiAP2 transcription factors that is involved in the regulation of gene expression during sexual development (e.g. Kafsack *et al*., 2014; Sinha *et al*., 2014; Poran *et al*., 2017; Bancells *et al*., 2019; van Biljon *et al*., 2019; Josling *et al*., 2020). Other ApiAP2 proteins of this group include AP2-G2 and AP2-G3, which co-regulate gametocytogenesis in conjunction with AP2-G (Yuda *et al*., 2015; Modrzynska *et al*., 2017; Zhang *et al*., 2017; Xu *et al*., 2021), and AP2-G5, which is essential for gametocyte maturation by down-regulation of *ap2-g* expression (Shang *et al*., 2021). Two other ApiAP2 domain transcription factors, AP2-FG and AP2-O3, contribute to sex differentiation. While AP2-FG binds to the promoters of many female-specific genes and is thus required for the establishment of the full female gene expression profile, AP2-O3 acts as a transcription repressor by targeting and repressing male genes in female gametocytes as a means of safeguarding the female-specific transcriptome (Yuda *et al*., 2020; Li *et al*., 2021).

In addition to transcriptional regulation by ApiAP2 proteins, post-transcriptional mechanisms are responsible for gametocyte formation and sex differentiation. Increasing evidence from recent years points to a critical role of RNA-binding proteins in the timely regulation of mRNA and protein synthesis during lifecycle progression of the parasite, including their involvement in gametocytogenesis (reviewed in e.g., Cui *et al*., 2015; Bennink and Pradel, 2019; Goyal *et al*., 2022). A series of RNA-binding proteins have been identified in *Plasmodium*, e.g. RNA helicases, zinc finger proteins (ZFPs; exhibiting C3H1 and C2H2 motifs), or members of the K homology (KH), Pumilio and Fem-3 binding factor (PUF) and acetylation lowers binding Affinity (ALBA) families (Reddy *et al*., 2015). While the function of most of these RNA-binding proteins is yet unknown, some proteins have been assigned to translationally regulating the intraerythrocytic replication cycle, e.g. the DNA/RNA binding protein ALBA1 and the two m^6^A-mRNA-binding YTH domain proteins YTH.1 and YTH.2 (Vembar *et al*., 2015; Baumgarten *et al*., 2019; Sinha *et al*., 2021). Others have been linked to stage conversion and transmission, e.g. ALBA4 with crucial functions in gametocytes and sporozoites of the rodent malaria parasite *P. yoelii* (Munoz *et al*., 2017), or PUF1 and PUF2 of *P. falciparum*, which are involved in gametocyte maturation and sex differentiation (Miao *et al*., 2010; Shrestha *et al*., 2016).

Of particular importance for human-to-mosquito transmission are RNA-binding proteins that act as translational repressors. Translational repression allows the gametocytes to rapidly react to external stimuli following their transmission to mosquitoes. The repressed transcripts are bound in messenger ribonucleoprotein particles (mRNPs), which condense to cytosolic aggregations. They are particularly present in female gametocytes, where they store mRNAs that encode proteins required for the development of the mosquito midgut stages and which are introduced to protein synthesis at the onset of gametogenesis (reviewed in e.g. Bennink and Pradel, 2019). A prominent repressor protein originally identified in female *P. berghei* gametocytes is DOZI (development of zygote inhibited; termed DZ50 in *P. falciparum*) and its interaction partner CITH (homolog of worm CAR-I and fly Trailer Hitch), which are responsible for repression of the transcripts encoding the zygote surface antigens P25 and P28 (Mair *et al*., 2006; 2010). A similar function has been assigned to PUF2 in *P. falciparum,* which also represses a number of female gametocyte transcripts including *p*25 and *p*28 (Miao *et al*., 2010, 2013).

A yet under-investigated group of RNA-binding proteins are ZFPs with C2H2 (CCHH) and C3H1 (CCCH) motifs. In particular, the C3H1-ZFPs commonly function in RNA-binding with important roles in pre-mRNA splicing, polyadenylation, mRNA export, and translation, as well as ubiquitination and transcriptional repression (reviewed in Ngwa *et al*., 2021). One of the few C3H1-ZFPs characterized so far is YTH.1, a binding protein of m^6^A-modified mRNA in early *P. falciparum* intraerythrocytic parasites that is crucial for post-transcription control (Baumgarten *et al*., 2019). A recent study further identified two *P. falciparum* C3H1-ZFPs, termed *Pf*CZIF1 and *Pf*CZIF2, suggested to assist in the regulation of the expression of proteins exported into the RBC cytosol after merozoite invasion (Balbin *et al*., 2023). In addition, a C3H1-ZFP of *P. falciparum* gametocytes, termed ZNF4, was assigned to male gametogenesis. ZNF4-deficient gametocytes were impaired in exflagellation, and comparative transcriptomics demonstrated the downregulation of male enriched genes associated to axonemal dynein complex formation, and cell projection organization in parasites lacking ZNF4 (Hanhsen *et al*., 2022).

The present study aims at investigating the role of the C3H1-ZFP MD3 (male development 3) in gametocyte development and gametogenesis. MD3 was previously identified by us during a transcriptomic screen for genes deregulated upon treatment of gametocytes with the histone deacetylase (HDAC) inhibitor, Trichostatin A (TSA) (Ngwa *et al*., 2017). Recently, MD3 was also identified as a protein involved in male gametocytogenesis in *P. berghei*, following a global screen of barcoded mutants (Russell *et al*., 2023). We here demonstrate that MD3 is highly expressed in male *P. falciparum* gametocytes and important for gametocyte development and exflagellation. Interactomics demonstrate its involvement in RNA-binding protein complexes which regulate the expression of sex-specific proteins.

## Results

### MD3 is a C3H1-ZFP predominantly expressed in immature male gametocytes

MD3 (PF3D7_0315600) is a 57-kDa protein with a C3H1-zinc finger domain spanning amino acid 163 to 185 (Fig. 1A). It was originally identified by us in a comparative transcriptomics screen for genes deregulated in immature gametocytes upon TSA treatment, suggesting its epigenetic regulation during gametocyte development (Ngwa *et al*., 2017). 3D structure prediction of MD3 demonstrated the arrangement of alpha helices and beta strands in the C3H1 domain to coordinate the zinc ion binding and domain stabilization (Fig. 1B, S1).

**Figure 1:**
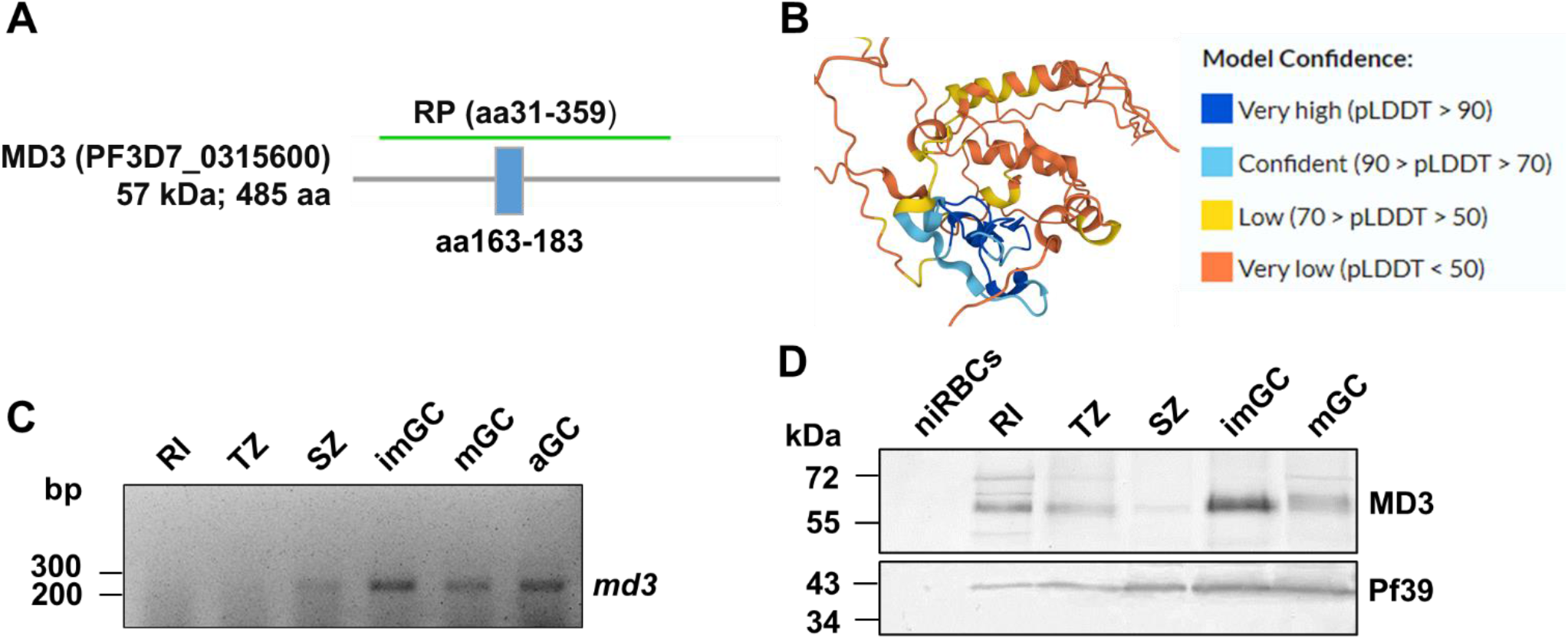
Structure of MD3 and expression in the *P. falciparum* blood stages. **(A)** Schematic depicting the MD3 protein. The region comprising the recombinant protein (RP) is indicated. Blue box, C3H1 domain. **(B)** Predicted 3D structure of the C3H1 domain of MD3. The 3D structure of the C3H1 zinc finger domain of MD3 was predicted, using the AlphaFold programme. Model confidence is shown by colour codes. The full 3D structure is provided in Fig. S1 **(C)** Transcript expression of MD3. Diagnostic RT-PCR was used to amplify *md3* transcript (231 bp) from cDNA generated from total RNA of rings (RI), trophozoites (TR), schizonts (SZ), as well as immature gametocytes (imGC; stages II-IV), mature gametocytes (mGC; stage V) and activated gametocytes (aGC; 30’ post-activation) of WT NF54. Controls are provided in Fig. S2A. **(D)** Protein expression of MD3. WB analysis of lysates from RI, TZ, SZ, imGC and mGC of WT NF54 using mouse anti-MD3 antisera detected MD3 with the expected molecular weight of ∼57 kDa. As a negative control, lysate of non-infected RBCs (niRBC) was used. Immunoblotting with rabbit antisera against *Pf*39 (∼39 kDa) served as loading control. Results (C, D) are representative of three independent experiments.

To study the expression profile of MD3, we first determined its transcript levels using diagnostic RT-PCR with RNA isolated from different asexual and sexual blood stages of the *P. falciparum* wildtype (WT) strain NF54. We detected high transcript abundance in immature, mature and activated gametocytes (30 min post-activation), while only low transcript levels were detected in rings, trophozoites or schizonts (Fig. 1C). Transcript analysis of the housekeeping gene *aldolase* was used as a loading control, and purity of the asexual blood stage and gametocyte samples was demonstrated by amplification of transcripts for the asexual blood stage-specific gene *ama1* (apical membrane antigen 1) and for the gametocyte-specific gene *ccp2* (LCCL-domain protein 2). The cDNA preparations lacking the reverse transcriptase were used to verify that samples were devoid of gDNA (Fig. S2A). These data confirmed transcription of *md3* in gametocytes and are in accord with previous studies (Lopez-Barragan *et al*., 2011; Ngwa *et al*., 2017).

We then generated mouse anti-MD3 polyclonal antisera against an N-terminal portion of MD3 for protein level analyses (Fig. 1A). Lysates were prepared from rings, trophozoites, schizonts, immature and mature gametocytes and subjected to Western blot (WB) analysis using the anti-MD3 antisera. Immunoblotting detected MD3 migrating at the expected molecular weight of 57 kDa in the asexual blood stages and gametocytes with high protein levels observed in immature gametocytes (Fig. 1D). Antisera against the endoplasmic reticulum-resident protein *Pf*39 were used as loading control, while lysate from non-infected RBCs served as negative control in the WB experiments.

Indirect immunofluorescence assays (IFAs) were employed to determine the subcellular localization of MD3 in the *P. falciparum* blood stages and revealed a granular expression of MD3 in the cytoplasm of all stages. MD3 was detected with high protein abundance in immature (stage II and III) gametocytes (Fig. 2A). While MD3-positive granules were evenly distributed in rings, trophozoites, and schizonts (highlighted by rabbit anti-P92 antisera), in gametocytes (highlighted by rabbit anti-P230 antisera), they were located partially near the periphery. In mature and activated gametocytes (30 min post-activation), the protein levels decreased. For negative control, schizont and gametocytes were immunolabelled with serum from non-immunized mice (Fig. S2B).

**Figure 2:**
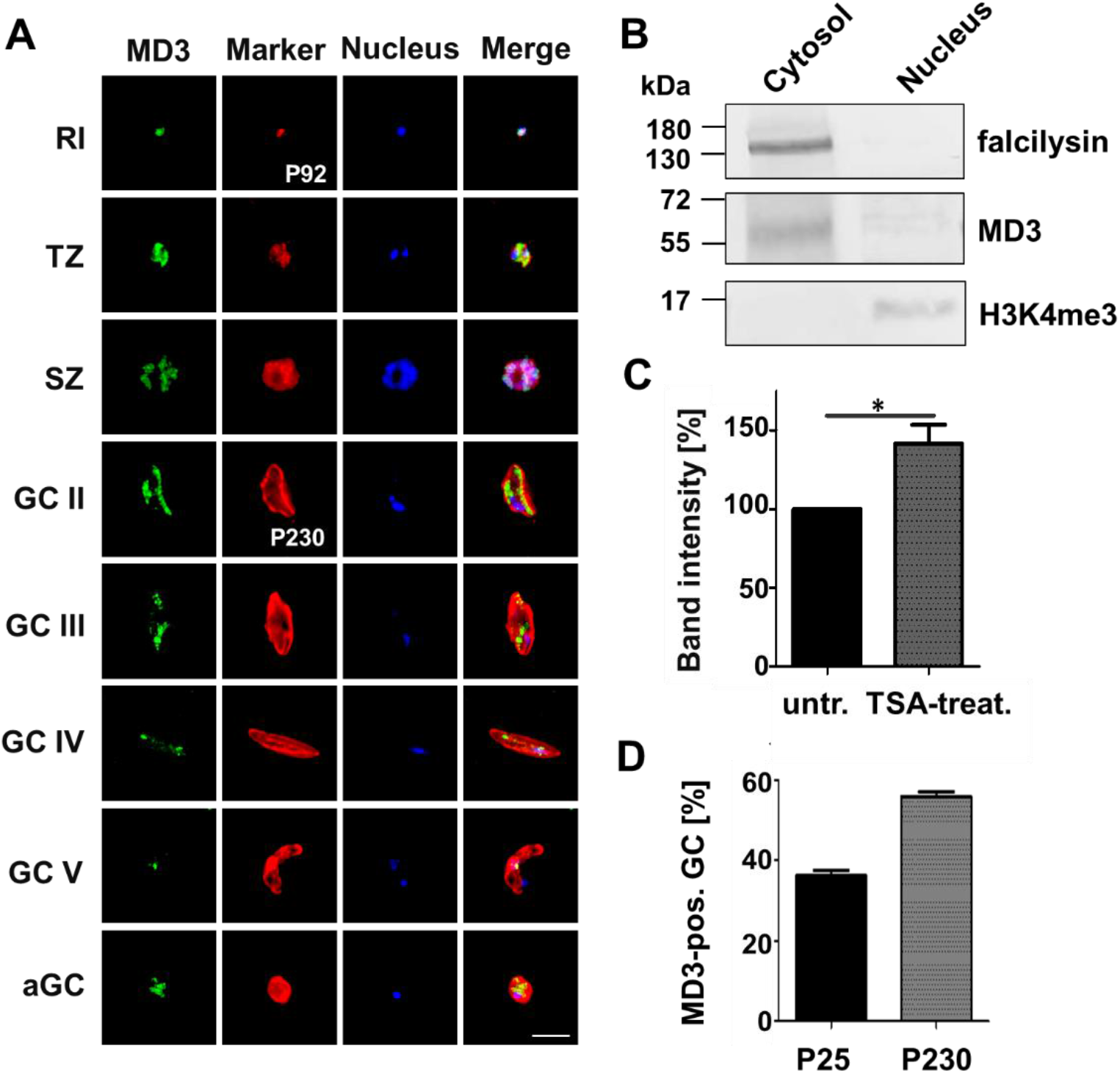
Subcellular localization, epigenetic regulation and sex specificity of MD3 in blood stage parasites. **(A)** Localization of MD3 in the *P. falciparum* blood stages. Methanol-fixed rings (RI), trophozoites (TZ), schizonts (SZ), gametocyte (GC) stages of II-V and activated gametocytes (aGC; 30 min post-activation) were immunolabeled with mouse anti-MD3 antisera (green). Asexual blood stages and gametocytes were highlighted by with rabbit antisera directed against P92 and P230, respectively (red); nuclei were highlighted by Hoechst 33342 nuclear stain (blue). Bar, 5 μm. Negative controls are provided in Fig. S2B. **(B)** Subcellular localization of MD3. Cytosolic and nuclear fractions were extracted from the immature gametocytes of WT NF54 and immunoblotted with mouse anti-MD3 antisera to detect MD3 (∼57 kDa). Mouse antisera against the cytosolic protease falcilysin (∼138 kDa) and rabbit antisera directed against the histone H3 mark H3K4me3 (∼15 kDa) were used to confirm the fraction purity. **(C)** Quantification of MD3 levels following TSA-treatment of immature gametocytes. Immature gametocytes were treated with 0.26 μM TSA or 0.5% (v/v) ethanol for control for 24 h. Lysates were subjected to WB, using anti-MD3 antisera. MD3 protein levels were evaluated by measuring the band intensities in four independent WB experiments using Image J; the values were normalized to the respective *Pf*39 protein band, used as loading control. Results are shown as mean ± SD (untreated set to 100%). *, p < 0.05 (Student’s t-test). An exemplary WB image is provided in Fig. S2C. **(D)** Sex specificity of MD3. WT NF54 mature gametocytes were immunolabeled with mouse anti-MD3 antisera and counterlabelled with either rabbit antisera against P230 (expressed in male and female gametocytes) or P25 (expressed in female gametocytes). For each marker, 100 labelled gametocytes were counted and the percentage of MD3-positive gametocytes was calculated. The experiment was performed in triplicate (mean ± SD). Results (A, B) are representative of three independent experiments.

Sub-cellular fractionation was conducted using lysates of immature gametocytes to obtain cytosolic and nuclear fractions, which were subjected to WB. MD3 was predominantly found in the cytosolic fraction, using the respective mouse anti-MD3 sera, while only minor signals were found in the nuclear fraction (Fig. 2B). Mouse antisera against the cytosolic protease falcilysin was used as a marker for the cytosolic fraction, while rabbit antibodies against the histone 3 methylation mark H3K4me3 served to confirm the nuclear fraction.

Because MD3 was originally identified by us as a product of a gene transcriptionally upregulated in immature gametocytes treated with the epigenetic HDAC inhibitor TSA (Ngwa *et al*., 2017; see above), we aimed to determine the protein levels of MD3 in dependence of TSA treatment. Lysates of immature gametocytes treated with 0.26 µM TSA for 24 h were harvested and immunoblotted with mouse anti-MD3 antisera, lysates of untreated cultures served as control. Quantitative WB demonstrated a significant increase of MD3 levels to 141% following TSA-treatment compared to untreated controls (set to 100%) in four independent experiments (Fig. 2C, S2C). These data confirm our previous findings that MD3 expression increases upon HDAC inhibition.

Potential sex specificity was determined by IFA as described previously (Bennink *et al*., 2018). MD3 was counterlabelled either with rabbit antisera against P230, a protein expressed in both female and male gametocytes, or with rabbit antisera against the female specific protein P25, and the numbers of MD3-positive gametocytes, positive for either P230 or P25 were determined. Noteworthy*, P. falciparum* gametocytes usually exhibit a male to female sex ratio of 1:4 (Smith *et al*., 2000). We showed that approximately 55% of total gametocytes (i.e. P230-positive gametocytes) labeled for MD3, while only 37% of female gametocytes (i.e. P25-positive gametocytes) were MD3-positive (Fig. 2D). The results indicate the presence of MD3 in both male and female gametocytes with a higher abundance in males and are in accord with previous reports on a high *md3* transcript levels in male gametocytes of *P. falciparum* and *P. berghei* (Lasonder *et al*., 2016; Russell *et al*., 2023).

The combined data indicate that MD3 is a cytosolic C3H1-ZFP particularly present in male immature gametocytes, the expression of which is dependent on HDAC-mediated epigenetic regulation.

### MD3 deficiency results in impaired intraerythrocytic growth, delayed gametocyte maturation and aborted exflagellation

For functional characterization of MD3, we employed the selected linked integration method using the pSLI-HA-*glmS* vector to generate line MD3-HA-*glmS* (Fig. S3A; Prommana *et al*., 2013; Birnbaum *et al*., 2017; Musabyimana *et al*., 2022), which expresses the protein of interest fused to a hemagglutinin A (HA)-tag. Further, the transcript contains the sequence for the *glmS* ribozyme in the 3’ untranslated region, enabling transcript degradation and hence conditional *md3-ha* knockdown upon addition of glucosamine (GlcN). Successful vector integration into the targeted *md3* locus was shown by diagnostic PCR (Fig. S3B). The MD3-HA-*glmS* line was free of wild type parasites.

First, the MD3-HA-*glmS* line was used to confirm protein expression in blood stage parasites. WB analysis using rat anti-HA antibody demonstrated expression of MD3-HA in the asexual and sexual blood stages with peak expression in immature gametocytes (Fig. S4A). No labeling was detected in non-infected RBCs and in mixed asexual cultures of the WT NF54 strain. Immunoblotting with rabbit antisera against *Pf*39 was used as a loading control. When lysates from immature gametocytes following subcellular fractionation were employed to WB, MD3-HA was primarily detected in the cytosol, while a minor band was additionally seen in the nuclear fraction (Fig. S4B). Immunoblotting with mouse anti-falcilysin antisera and rabbit anti-H3K4me3 antibody were used as cytosolic and nuclear fraction controls as described before. In addition, MD3-HA expression in immature gametocytes was upregulated to 135% (untreated set to 100%), when these were treated with TSA as described above (Fig. S4C, D).

IFAs further confirmed the granular localization of MD3-HA in the cytoplasm of asexual blood stages and gametocytes of line MD3-HA-*glmS* (highlighted by Evans blue and rabbit anti-P230 labeling, respectively) while no MD3-HA labeling was detected in the WT NF54 blood stages (Fig. S5A). MD3-HA was predominantly found in male gametocytes, when counterlabelling experiments were performed. Approximately 59% of total gametocytes (i.e. P230-positive gametocytes) and 43% of female gametocytes (i.e. P25-positive gametocytes) were MD3-HA-positive (Fig. S5B, C).

Subsequently, the MD3-HA-*glmS* line was used for loss-of-function phenotyping. Treatment of the MD3-HA-*glmS* asexual blood stages with 5 mM GlcN for 72 h reduced the MD3-HA protein levels to 63% compared to untreated cultures (set to 100%), as demonstrated by quantitative WB (Fig. S6A, B). In the following, tightly synchronized ring stages of line MD3-HA-*glmS* with an initial parasitemia of 0.25% were cultivated with 5 mM GlcN and the parasitemia was followed up by Giemsa-stained thin blood smears every 24 h over a period of 96 h. Untreated cultures of line MD3-HA-*glmS* as well as GlcN-treated and untreated WT NF54 were carried along for controls. The intraerythrocytic growth assay revealed a significant reduction in the parasitemia of GlcN-treated MD3-HA-*glmS* at 96 h as compared to the controls, indicating that the knockdown of *md3-ha* transcript impairs asexual blood stage development (Fig. 3A). However, no morphological abnormalities were observed in the GlcN-treated blood stage parasites compared to the untreated transgenic line or GlcN-treated and untreated WT NF54, as demonstrated by Giemsa smears (Fig. S6C). Further, no stage conversion shifts were recorded, when the numbers of ring stages, trophozoites and schizonts were evaluated in the four different experimental setups (Fig. S7A).

**Figure 3:**
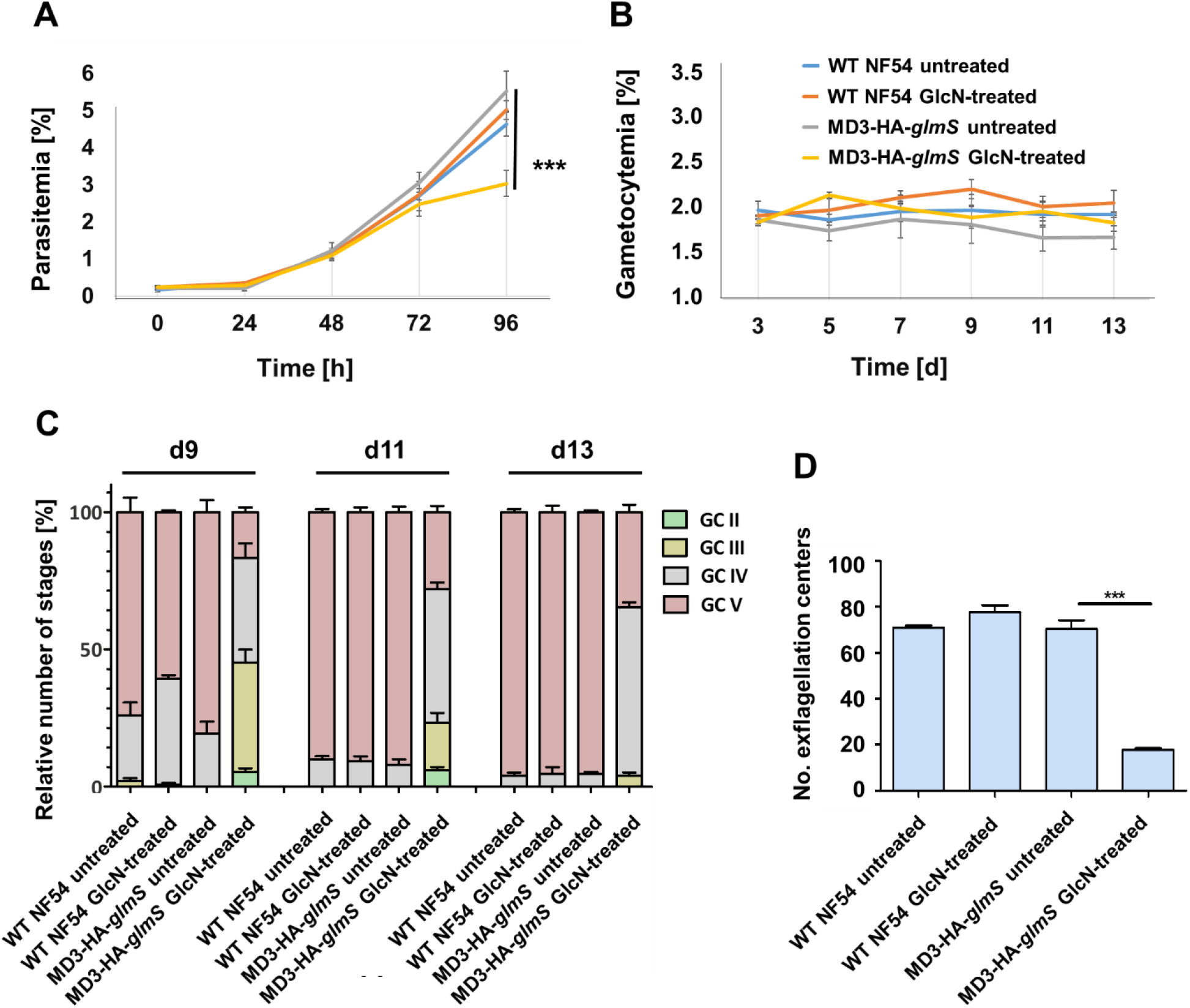
The effect of MD3-HA knockdown on intraerythrocytic growth and gametocyte development. **(A)** Asexual blood stage replication following MD3-HA knockdown. Synchronized ring stage cultures of WT NF54 and line MD3-HA-*glmS* with a starting parasitaemia of 0.25% were maintained in cell culture medium supplemented or not with 5 mM GlcN for transcript knockdown. The parasitaemia was followed via Giemsa smears over a time-period of 0 - 96 h. The experiment was performed in triplicate (mean ± SD). ***, p < 0.001 (One-way ANOVA with post hoc Bonferroni multiple comparison test). **(B)** Gametocyte development following MD3-HA knockdown. Gametocyte production was induced in synchronized ring stage cultures of WT NF54 and line MD3-HA-*glmS* at a final parasitaemia of 7.5% by addition of lysed RBCs. Subsequently, the cultures were maintained in cell culture medium supplemented or not with 5 mM GlcN for transcript knockdown and the gametocytaemia was followed via Giemsa smears every 48 h between day 3 – 13 post-induction. The experiment was performed in triplicate (mean ± SD). **(C)** Gametocyte maturation following MD3-HA knockdown. Relative numbers of gametocyte stages II-V were followed via Giemsa smears in 50 iRBCs over a time-period of 13 d. The experiment was performed in triplicate. The complete assay is shown in Fig. S7B. **(D)** Effect of MD3-HA knockdown on exflagellation. Gametocyte production was induced in synchronized ring stage cultures of WT NF54 and line MD3-HA-*glmS* at a final parasitaemia of 7.5% by addition of lysed RBCs. Following treatment with 20 U/ml of heparin for 2 days, cultures were maintained in normal cell culture medium. On day 7-10 post-induction, the gametocytes were maintained in cell culture medium supplemented or not with 5 mM GlcN for transcript knockdown On day 13, exflagellation centers were counted after activation with 100 µM xanthurenic acid (XA) for 15 min at RT. The experiment was performed in triplicate (mean ± SD). ***, p < 0.001 (One-way ANOVA with post hoc Bonfferoni multiple comparison test). Exemplary images of mature gametocytes used in the exflagellation assays are provided in Fig. S8.

In order to determine the effect of MD3-HA deficiency on gametocytogenesis, tightly synchronized ring stage parasite cultures of line MD3-HA-*glmS* at a final parasitemia of 7.5% were induced to form gametocytes by addition of lysed RBCs for 24 h. After induction, cultures were maintained in cell culture medium supplemented with 20 U/ml heparin to kill the asexual blood stages for 4 days and with 5 mM GlcN for transcript knockdown till day 13. Samples were taken every 48 h starting from day 3 after induction and subjected to Giemsa smear staining. The gametocytemia was evaluated and the numbers of stage II to V gametocytes were determined microscopically. The gametocytemia was not affected in the GlcN-treated MD3-HA-*glmS* line as compared to untreated transgenic parasites or GlcN-treated and untreated WT NF54 (Fig. 3B). However, a delay in gametocyte maturation was observed in the MD3-HA-*glmS* line at day 13 of GlcN treatment compared to controls. Here, the majority of gametocytes were detected at stage IV, while the control parasites have progressed to stage V (Fig. 3C, S7B). No morphological differences were observed in the GlcN-treated MD3-HA-*glmS* gametocytes compared to the controls, though, as shown by Giemsa staining (Fig. S6C).

Finally, the effect of MD3 deficiency was studied during male gametogenesis. Highly synchronized ring stage cultures of line MD3-HA-*glmS* were induced to form gametocytes. After 24 h, the asexual blood stages were killed by treatment with heparin for 2 days as described above. At day 7 post-induction, the MD3-HA-*glmS* gametocytes were treated with 5 mM GlcN for 4 days. The gametocytes were then allowed to recover for 2 days. Untreated cultures of line MD3-HA-*glmS* as well as GlcN-treated and untreated WT NF54 were carried along for control. Exflagellation of the male gametocytes was determined on day 13 by microscopy. For this, the gametocyte cultures of the four setups were adjusted to total RBC numbers and the numbers of mature gametocytes were determined for each culture by Giemsa staining (Fig. S8). Following gametocyte activation, the number of exflagellation centers were counted and the results were adjusted to the numbers of mature gametocytes as evaluated prior to activation. A significant reduction in the number of exflagellation centers was observed for line MD3-HA-*glmS* treated with GlcN compared to the controls (Fig 3D). The combined data demonstrate that MD3 deficiency impairs male gametocyte development and in consequence gametogenesis.

The combined phenotyping data indicate MD3 is crucial for both intraerythrocytic development and gametocyte maturation with a significant effect on male gametogenesis.

### The MD3 interaction network is composed of regulators of sexual development

To determine the MD3 interaction network, we generated transgenic lines episomally expressing a MD3-GFP-BirA fusion protein as described (Musabyimana *et al*., 2022). Blood stage parasites were transfected with the vectors pARL-MD3-*pfama1*-GFP-BirA or pARL-MD3-*pffnpa*-GFP-BirA, whereby the expression of MD3-GFP-BirA was either controlled by the asexual blood stage-specific *pfama1* or the gametocyte-specific *pffnpa* promotor (Fig. S9A, B). Diagnostic PCR confirmed the presence of the respective vectors in the transgenic lines (Fig. S9C, D).

WB analysis, using mouse anti-GFP antibody, demonstrated the expression of the MD3-GFP-BirA fusion protein in lysates from asexual blood stages and gametocytes of lines MD3-*pfama1*-GFP-BirA and MD3-*pffnpa*-GFP-BirA, respectively. In the transgenic lines, MD3-GFP-BirA could be detected at the expected molecular weight of approximately 118 kDa as compared to WT NF54 parasites, where no signal was present (Fig. 4A). In addition, immunolabeling with anti-GFP antibody confirmed the cytoplasmic localization of MD3-GFP-BirA in trophozoites and gametocytes (highlighted by rabbit anti-Pf39 and P230 antisera, respectively) of the respective lines, while no signal was detected in the WT NF54 control (Fig. 4B).

**Figure 4:**
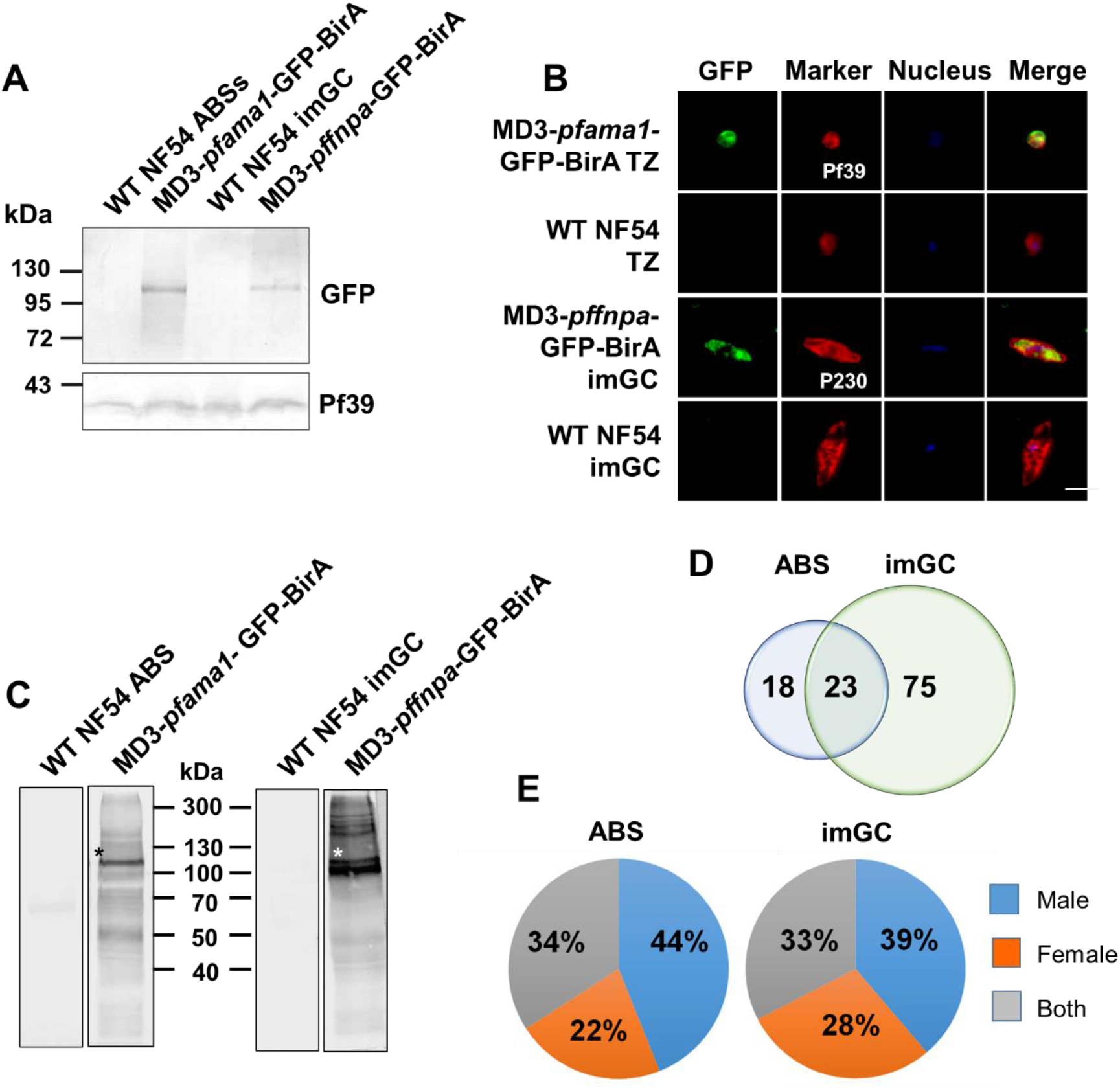
Verification of the MD3-GFP-BirA parasite lines to be used in BioID. **(A)** Verification of MD3-GFP-BirA expression. Lysates of mixed asexual blood stages (ABS) of line MD3-*pfama1*-GFP-BirA and purified immature gametocytes (imGC) of line MD3-*pffnpa*-GFP-BirA were subjected to WB analysis using mouse anti-GFP antibody to detect MD3-GFP-BirA running at the expected molecular weight of 118 kDa. Lysates of the respective WT NF54 stages were used for negative controls. Rabbit antisera against *Pf*39 (∼39 kDa) was used as a loading control. **(B)** Localization of MD3-GFP-BirA in blood stage parasites. Methanol-fixed trophozoites (TZ) of line MD3-*pfama1*-GFP-BirA and immature gametocytes (imGC) of line MD3-*pffnpa*-GFP-BirA were immunolabeled with mouse anti-GFP antibody to detect MD3-GFP-BirA (green). Trophozoites were counterlabelled with rabbit anti-*Pf*39 antibody and gametocytes with rabbit anti-P230 antisera (red); nuclei were highlighted by Hoechst 33342 nuclear stain (blue). WT NF54 was used as a negative control. Bar, 5 μm. **(C)** Detection of biotinylated proteins in the MD3-GFP-BirA lines. Asexual blood stages (ABS) of line MD3-*pfama1*-GFP-BirA and immature gametocytes (imGC) of line MD3-*pffnpa*-GFP-BirA were treated with 50 μM biotin for 24 h. Lysates were subjected to WB analysis and protein biotinylation was detected using streptavidin coupled to alkaline phosphatase. Asterisks indicate biotinylated MD3. Biotin-treated WT NF54 was used as a negative control. **(D)** Venn diagram depicting the MD3 interactors grouped by blood stage. Asexual blood stages (ABS) and immature gametocytes (imGC) were treated with biotin as described above and streptavidin bead-purified biotinylated proteins were subjected to BioID-MS. Hits with predicted signal peptides were excluded from further analysis. A total of 116 interactors were identified and grouped by blood stage. **(E)** Pie chart depicting the MD3 interactors (percentage of total numbers) grouped by sex (see PlasmoDB database). Results (A-C) are representative of three independent experiments. Detailed information on the interactors is provided in Tables S1 and S2.

Subsequently, lysates were prepared from asexual blood stages and gametocytes of lines MD3-*pfama1*-GFP-BirA and MD3-*pffnpa*-GFP-BirA, respectively, which had been incubated with 50 μM biotin for 24 h. Immunoblotting, using alkaline phosphatase-conjugated streptavidin, led to the detection of multiple protein bands in both lysates, indicative of biotinylated proteins, including a protein of approximately 118 kDa representing biotinylated MD3 (Fig. 4C). No biotin-positive protein bands were detected in WT NF54. Furthermore, immunolabeling with fluorophore-conjugated streptavidin showed the presence of biotinylated proteins in the cytoplasm of biotin-treated trophozoites and gametocytes of the respective transgenic lines (highlighted by rabbit anti-Pf39 and P230 antisera, respectively), while no biotinylated proteins were detected in the WT NF54 (Fig. S9E).

Lines MD3-*pfama1*-GFP-BirA and MD3-*pffnpa*-GFP-BirA were subjected to mass spectrometry-based proximity-dependent biotin identification (BioID-MS) to identify the MD3 interactome. Schizonts and immature gametocytes of the respective lines were treated with 50 µM biotin for 24 h and equal amounts of parasites were harvested. Three independent samples were collected from each of the two lines and mass spectrometric analysis was performed on streptavidin-purified protein samples with three technical replicas for each sample. Pull-down samples from WT NF54 lysates were used as a negative control. BioID-MS resulted in the identification of 41 significantly enriched hits in the asexual blood stages of line MD3-*pfama1*-GFP-BirA (after exclusion of proteins with a signal peptide) and 98 significantly enriched hits in gametocytes of line MD3-*pffnpa*-GFP-BirA compared to WT NF54 control (Fig. 4D; Table S1, S2). A total of 23 proteins were found in both lines, which included the bait protein MD3. *In-silico* transcript analyses (according to table “Transcriptomes of 7 sexual and asexual life stages”; López-Barragán *et al.,* 2011; see PlasmoDB database; Aurrecoechea *et al.,* 2009) showed that the majority of 98 MD3 interactors identified in line MD3-*pffnpa*-GFP-BirA exhibited peak expression in ookinetes and mature gametocytes in addition to ring stages and trophozoites, and most of the 41 MD3 interactors identified in the MD3-*pfama1*-GFP-BirA line had peak expression in ring stages and early trophozoites (Fig. S10). When the sex specificity of the interactors was evaluated (according to table “Gametocyte Transcriptomes”; Lasonder *et al.,* 2016; see PlasmoDB database; Aurrecoechea *et al.,* 2009), roughly 40% of the MD3 interactors of immature gametocytes and asexual blood stages could be assigned to the male sex (Fig. 4E).

Gene Ontology (GO) enrichment analysis of the 98 MD3 interactors of immature gametocytes, using ShinyGO 0.77, revealed their association with biological processes such as translation, post-transcriptional regulation of gene expression and macromolecule metabolic processes (Fig. 5A). The interactors were primarily assigned to the cytoplasm and had associations with ribosomes and mRNP complex, e.g., eukaryotic translation initiation factor 4F complex, CCR4-NOT complex and P-bodies (Fig. 5B). Molecular functions included mRNA- and rRNA-binding as well as binding to heterocyclic and organic cyclic compounds (Fig. 5C). The 41 interactors of asexual blood stages were assigned to biological processes such as translation and gene expression with cellular component associations to the cytoplasm and RNP complexes, while mRNA-binding, rRNA-binding as well as organic cyclic compound binding represented the enriched terms for molecular functions (Fig. S11A-C).

**Figure 5:**
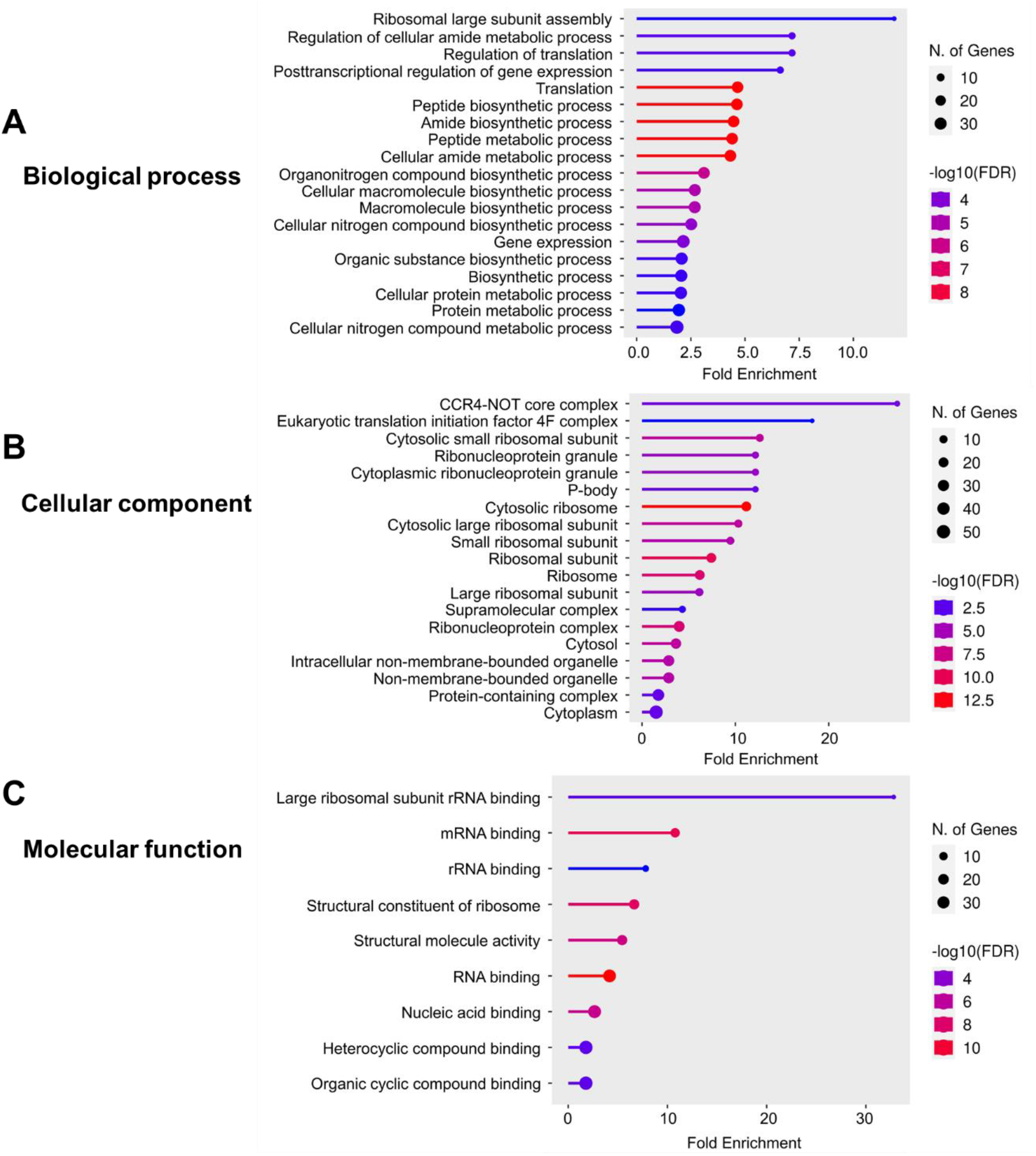
Functional prediction analysis of the MD3 interactors in immature gametocytes. A GO enrichment analysis of the 98 putative MD3 interactors in immature gametocytes was performed using the ShinyGO 0.77 programme with p < 0.05 to display the enriched GO terms based on biological process **(A)**, cellular component **(B)** and molecular function **(C)**.

The putative MD3 interactors were further subjected to STRING-based analyses to investigate the protein-protein interaction networks (see string-db.org; text mining included). The MD3 interactome of immature gametocytes comprised four defined clusters, the most comprehensive of which included multiple ribosomal proteins (Fig. 6, cluster A; Table S3; https://string-db.org/cgi/network?taskId=bdYMSO1hLfe9&sessionId=bklS63OmyZ9h). A similar cluster was identified for the MD3 interactors in the asexual blood stages (Fig. S12; Table S4; https://string-

**Figure 6:**
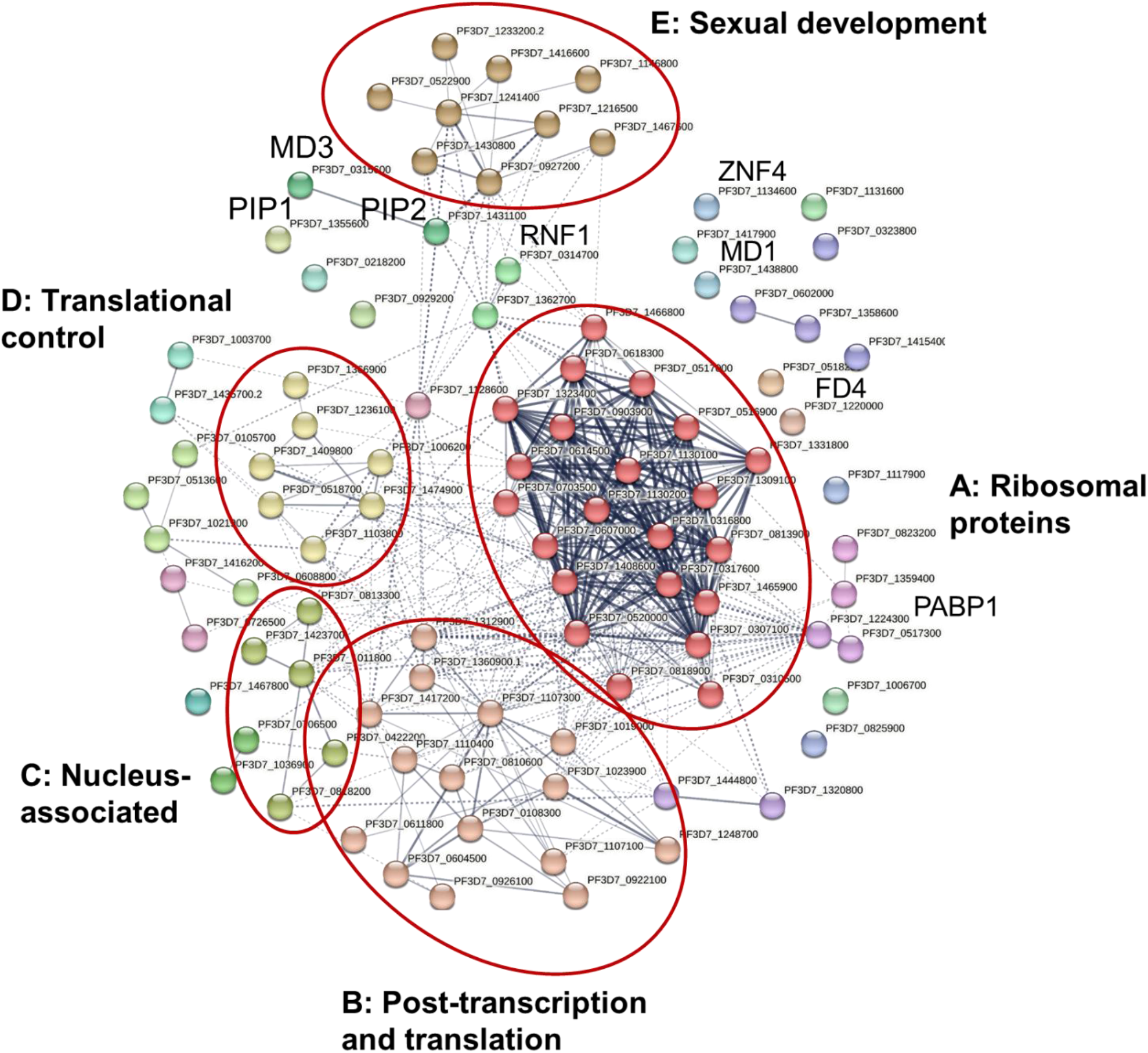
Network analysis of the MD3 interactors in immature gametocytes. A protein-protein network analysis of the 98 putative MD3 interactors in immature gametocytes was performed, using the STRING database, selecting a medium interaction confidence of 0.4. Line thickness specifies the strength of data support between the proteins. Clustering of the interactors was employed by the Markov Clustering (MCL) algorithm with an inflation parameter of 3; dotted lines represent the connection among the hits of different clusters. On the basis of physical interactions among the query proteins, five main clusters were identified, i.e. A) ribosomal proteins B) post-transcription and translation C) nucleus-associated D) translational control, and E) sexual development. Detailed information on each cluster and its components is provided in Table S3.

<u>db.org/cgi/network?taskId=baQUg0cZ9zAa&sessionId=b8m6z09Rup1D</u>). Three more clusters of MD3 interactors in immature gametocytes (clusters B-D) were tightly linked to processes of transcription and translation. Cluster B comprised 15 proteins associated to post-transcription and translation, e.g. NOT family proteins, eukaryotic translation initiation factors or RNA-binding proteins, while cluster C included five proteins predicted to be located to the nucleus, including a NPL domain protein and a NTF2 domain protein of unknown functions (Fig. 6; Table S3). Cluster D was formed by seven proteins linked to translational control, e.g. the RNA-binding proteins PUF1 and ALBA3 or the translational repressor CITH (Mair *et al*., 2010; Bunnik *et al*., 2016; Shresta *et al*., 2016; Banerjee *et al*., 2023). The most interesting cluster of the STRING analysis was cluster E that comprised nine proteins as well as various satellite proteins. The majority of these proteins had recently been linked to the development of male and female gametocytes, e.g. the cluster proteins GD1, FD1, FD2, MD2, MDV1 and the satellite proteins FD4, MD1, MD3, ZNF4 and RNF1 (Furuya *et al*., 2005; Ngwa *et al*., 2017; Gomes *et al*., 2022; Hanhsen *et al*., 2022; Russell *et al*., 2023).

The combined BioID-MS results unveiled an important interaction network of MD3 with proteins related to post-transcriptional and translational control in immature gametocytes and highlighted a striking interaction of MD3 with other RNA-binding proteins, including various ZFPs, involved in the regulation of gametocyte development.

### MD3 forms a multi-protein complex with RNA-binding proteins

In a final set of experiments, we investigated in more detail the interaction of MD3 and select RNA-binding proteins as identified by BioID-MS, using co-immunoprecipitation assays. We made use of the previously published ZNF4-HA-*glmS* line (Hanhsen *et al*., 2022). In addition, we generated a parasite line expressing PUF1 fused to an HA-tag, using the pSLI-HA-*glmS* vector as described above. Successful vector integration into the targeted *puf1* locus was shown by diagnostic PCR (Fig. S13A). Expression of HA-tagged PUF1 was subsequently shown WB analysis using rat anti-HA antibody. Immunoblotting highlighted a protein band running at the expected molecular weight of ∼227 kDa in the lysates of asexual blood stages and immature gametocytes of PUF1-HA-*glmS* parasites, while no band was visible in the WT NF54 control (Fig. S12B).

For the co-immunoprecipitation assays, lysates were generated from immature gametocytes of lines MD3-HA-*glmS*, PUF1-HA-*glmS*, and ZNF4-HA-*glmS*; WT NF54 lysate was used as a control. The bait proteins were precipitated with protein-G bead-bound rabbit anti-HA or rat anti-HA antibodies and precipitated proteins were subjected to WB analysis. In immunoprecipitates of line MD3-HA-*glmS*, protein bands representing RNF1, CITH and the polyadenylate-binding protein PABP1 could be detected running at the expected molecular weights, following immunoblotting with the respective mouse or rabbit antibodies (Fig. 7A, B). When immunoprecipitates of lines ZNF4-HA-*glmS* and PUF1-HA-*glmS* were subjected to WB analysis, MD3 could be detected using the respective mouse antibody (Fig. 7C, D). The precipitation of the respective bait protein was confirmed by immunoblotting with rat anti-HA antibody; immunoblotting with rabbit anti-*Pf*39 antibody was used as a negative control. In addition, WT NF54 was subjected to co-immunoprecipitation assays and no bands for precipitated parasite proteins were visible (Fig. S14A-D).

**Figure 7:**
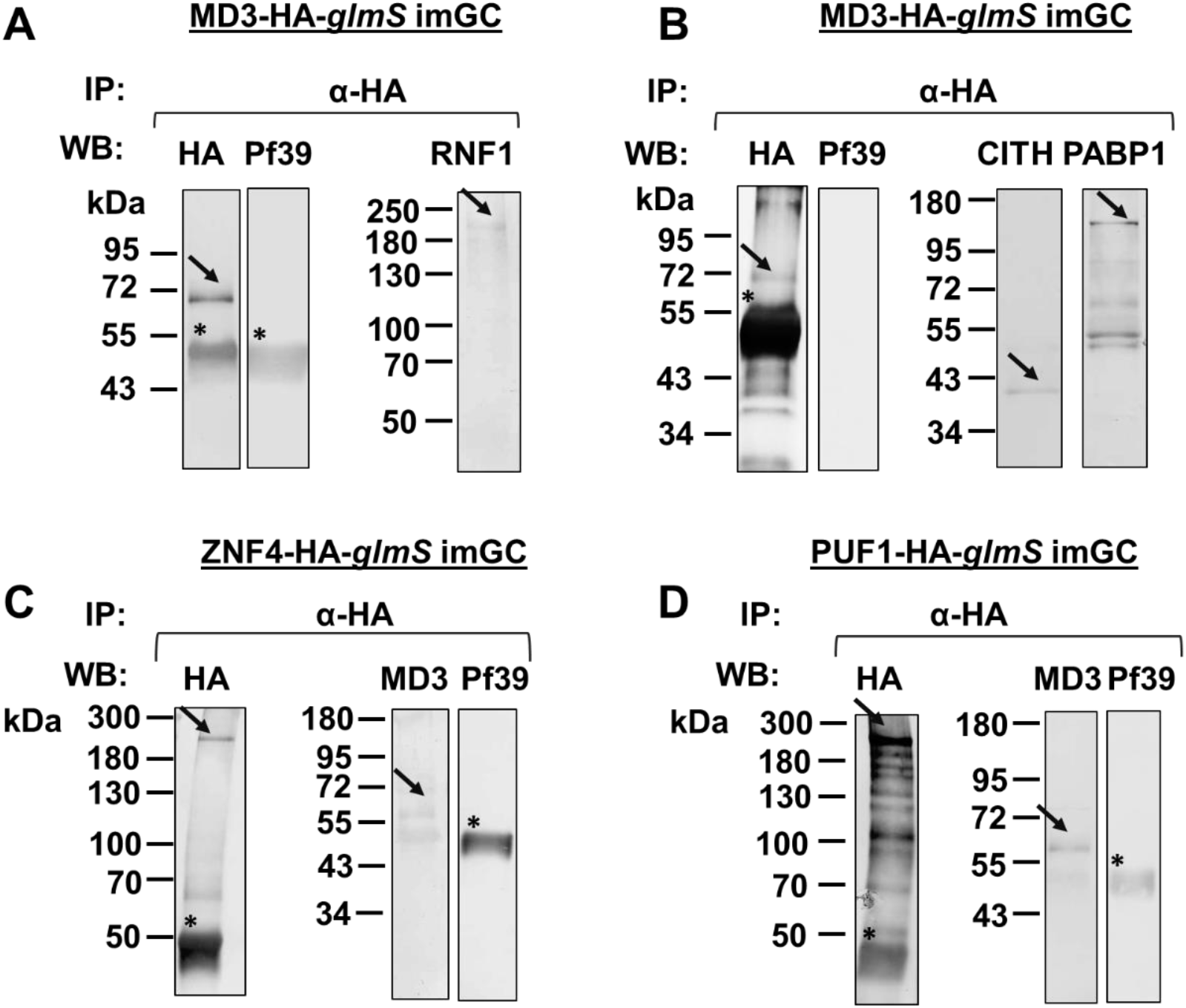
Protein interactions of MD3 with RNA-binding proteins. Lysates of immature gametocytes (imGC) of lines MD3-HA-*glmS* **(A, B)**, ZNF4-HA-*glmS* **(C),** and PUF1-HA-*glmS* **(D)** were subjected to co-immunoprecipitation assays. Protein complexes were immunoprecipitated using polyclonal rabbit or rat anti-HA antibodies, followed by immunoblotting using rabbit or rat anti-HA antibodies, mouse anti-RNF1 antisera, rabbit anti-CITH antibody, rabbit anti-PABP1 antibody or mouse anti-MD3 antisera to detect the precipitated proteins (expected molecular weights: MD3-HA, ∼60 kDa; ZNF4-HA, ∼207 kDa, PUF1-HA, ∼227 kDa; RNF1, ∼138 kDa, MD3, ∼57 kDa, CITH, ∼39 kDa, PABP1, ∼97 kDa). Arrows indicate the precipitated proteins. Immunoblotting with rabbit anti-*Pf*39 served as a negative control (∼39 kDa). Asterisks signify bands corresponding to the heavy chains of the precipitation antibody. Results are representative of two to three independent experiments.

In conclusion, the co-immunoprecipitation data confirmed protein-protein interactions between MD3 and selected RNA-binding proteins of immature gametocytes.

## Discussion

Gametocytogenesis of malaria parasites requires a tight control of transcriptional and translational regulation and RNA-binding proteins play crucial role during this process. In this study, we report on the importance of the C3H1-ZFP MD3 in regulating male gametogenesis by interacting with RNA-binding proteins. We show that MD3 is expressed in the blood stages of *P. falciparum* mainly present in the cytosol of immature male gametocytes, where it can be found in granular structures. We also confirmed the dependence of MD3 expression on HDAC-mediated epigenetic regulation with inhibition of HDAC activity resulting in higher MD3 levels. Further, MD3 deficiencies affect intraerythrocytic replication and particular impair gametocyte maturation and exflagellation.

To unveil the MD3 interactome, we employed BioID-MS analyses and identified 41 putative interactors in the asexual blood stages and 98 interactors in immature gametocytes; 23 proteins were shared by both stages. In-depth bioinformatic analyses demonstrated a strong link of these interactors with various RNA-binding complexes like the CCR4-NOT complex, the eukaryotic translation initiation complex, mRNP complexes and P-bodies/stress granules in addition to ribosomal proteins suggesting the involvement of MD3 in the regulation of mRNA processing and translation (the most important interactors are summarized in Table 1). STRING analyses confirmed these interactions by highlighting four distinct clusters in immature gametocytes, i.e. the post-transcription and translation cluster (cluster B), a nucleus-associated protein cluster (cluster C), the translational control cluster (cluster D), and a cluster comprising RNA-binding proteins recently linked to regulating sexual development (cluster E), in addition to a ribosome-related cluster that was also found in the interactome of asexual blood stages (cluster A).

**Table 1.**
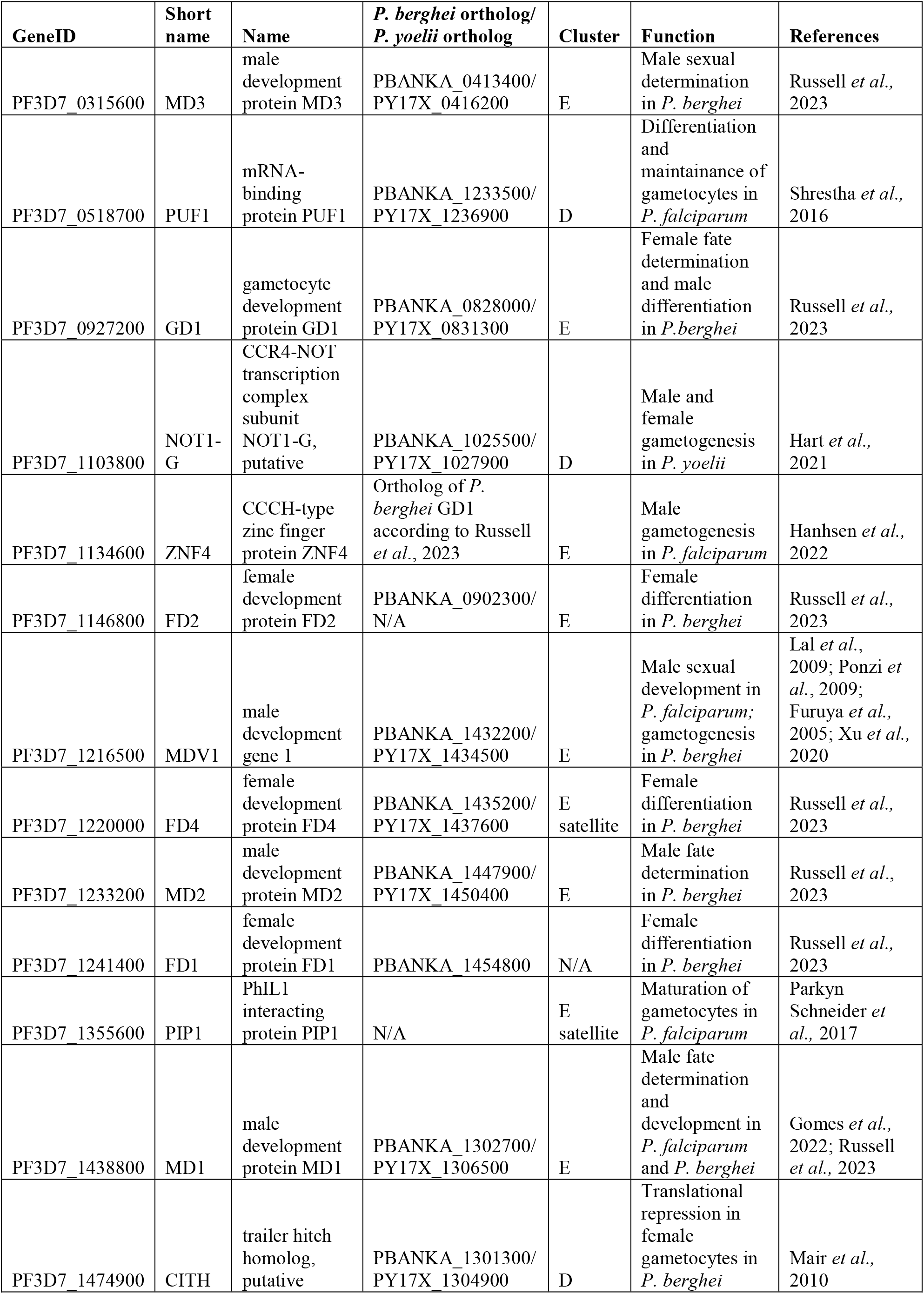
Selected interactors of *P. falciparum* MD3.

| GeneID | Short name | Name | *P. berghei* ortholog/<br>*P. yoelii* ortholog | Cluster | Function | References |
| --- | --- | --- | --- | --- | --- | --- |
| PF3D7_0315600 | MD3 | male development protein MD3 | PBANKA_0413400/<br>PY17X_0416200 | E | Male sexual determination in *P. berghei* | Russell *et al.*, 2023 |
| PF3D7_0518700 | PUF1 | mRNA-binding protein PUF1 | PBANKA_1233500/<br>PY17X_1236900 | D | Differentiation and maintainance of gametocytes in *P. falciparum* | Shrestha *et al.*, 2016 |
| PF3D7_0927200 | GD1 | gametocyte development protein GD1 | PBANKA_0828000/<br>PY17X_0831300 | E | Female fate determination and male differentiation in *P. berghei* | Russell *et al.*, 2023 |
| PF3D7_1103800 | NOT1-G | CCR4-NOT transcription complex subunit NOT1-G, putative | PBANKA_1025500/<br>PY17X_1027900 | D | Male and female gametogenesis in *P. yoelii* | Hart *et al.*, 2021 |
| PF3D7_1134600 | ZNF4 | CCCH-type zinc finger protein ZNF4 | Ortholog of *P. berghei* GD1 according to Russell *et al.*, 2023 | E | Male gametogenesis in *P. falciparum* | Hanhseen *et al.*, 2022 |
| PF3D7_1146800 | FD2 | female development protein FD2 | PBANKA_0902300/<br>N/A | E | Female differentiation in *P. berghei* | Russell *et al.*, 2023 |
| PF3D7_1216500 | MDV1 | male development gene 1 | PBANKA_1432200/<br>PY17X_1434500 | E | Male sexual development in *P. falciparum*; gametogenesis in *P. berghei* | Lal *et al.*, 2009; Ponzi *et al.*, 2009; Furuya *et al.*, 2005; Xu *et al.*, 2020 |
| PF3D7_1220000 | FD4 | female development protein FD4 | PBANKA_1435200/<br>PY17X_1437600 | E satellite | Female differentiation in *P. berghei* | Russell *et al.*, 2023 |
| PF3D7_1233200 | MD2 | male development protein MD2 | PBANKA_1447900/<br>PY17X_1450400 | E | Male fate determination in *P. berghei* | Russell *et al.*, 2023 |
| PF3D7_1241400 | FD1 | female development protein FD1 | PBANKA_1454800 | N/A | Female differentiation in *P. berghei* | Russell *et al.*, 2023 |
| PF3D7_1355600 | PIP1 | PhIL1 interacting protein PIP1 | N/A | E satellite | Maturation of gametocytes in *P. falciparum* | Parkyn Schneider *et al.*, 2017 |
| PF3D7_1438800 | MD1 | male development protein MD1 | PBANKA_1302700/<br>PY17X_1306500 | E | Male fate determination and development in *P. falciparum* and *P. berghei* | Gomes *et al.*, 2022; Russell *et al.*, 2023 |
| PF3D7_1474900 | CITH | trailer hitch homolog, putative | PBANKA_1301300/<br>PY17X_1304900 | D | Translational repression in female gametocytes in *P. berghei* | Mair *et al.*, 2010 |

Clusters A-D lie in close proximity to each other and are thematically linked. Here found are proteins of the CCR4-NOT and the eukaryotic translation initiation complexes as well as RNA-binding proteins. Some of the most prominent interactors of MD3 include known activators of translation, for example PABP, PAIP and eIF4G. For example, the translation initiation factor eIF4G (together with eIF4A and eIF4E) is linked to the mRNA 5’-cap and interacts with PABP and PAIP1 to facilitate mRNA pseudo-circularization and hence transcript stabilization. Interestingly, in *P. falciparum*, the DHH1/DDX6-like RNA helicase DOZI has been identified as a potential eIF4A protein and shown to interact with eIF4E (Tarique *et al*., 2013). Noteworthy, the PABP-mediated mRNA closed-loop can also facilitate transcript destabilization by providing the physical contact between mRNA-binding proteins, which typically target the 3′-UTR (untranslated region), and the 5′-UTR of the transcript in order to suppress translation (reviewed in Bennink and Pradel, 2019).

MD3 further interacts with components important for transcript decay, like the NOT proteins of the CCR4-NOT complex, a deadenylase complex that contributes to poly(A) tail-shortening. Importantly, one of the MD3 interactors, NOT1-G, has been described as a regulator of sexual stage maturation in *P. yoelii*. NOT1-G localizes to cytoplasmic granules in gametocytes and its deficiencies lead to defects in the formation of male gametes as well as of zygotes that derived from female gametes (Hart *et al*., 2021). Relevant to this, CCR4-1 was also reported to be crucial for the development and activation of male *P. yoelii* gametocytes as genetic disruption or catalytic inactivation of CCR4-1 resulted in defective male gametocyte maturation and hence, parasite transmission (Hart *et al*., 2019).

A variety of RNA-binding proteins mediate the interaction between the CCR4-NOT complex and the 3′-UTR, including PUF proteins, which can increase decapping and deadenylation, or the decapping DDH1/DDX6 activator that associates with Not1, and most of these factors are conserved in *P. falciparum* (reviewed in Bennink and Pradel, 2019). Importantly, some of the transcript decay factors were identified as interactors of MD3, like PUF1, which contributes to the maintenance and differentiation of *P. falciparum* gametocytes, with lack of PUF1 leading to a sharp decline in the late stage gametocytes and a sex-ratio shift to males (Shrestha *et al*.,,2016). In *P. falciparum*, two more members of the PUF family are known, PUF2 and PUF3. While PUF3 was reported to participate in ribosome biogenesis, *puf2* gene knockout promotes the formation particularly of male *P. falciparum* gametocytes (Miao *et al*., 2010). Moreover, PUF2 binds the transcripts of the zygote surface antigens P25 and P28 via PUF-binding elements in their 3′-UTR, thereby contributing to the translational repression of these transcript (Miao *et al*., 2013a).

Of particular importance is the interaction of MD3 with CITH, a master regulator of translational repression in female gametocytes as described earlier (reviewed in Bennink *et al*., 2016). CITH was previously shown to interact with DOZI and together they repress mRNA encoding P25 and P28 in female P*. berghei* gametocytes (Mair *et al*., 2006; 2010). Characterization of DOZI- and CITH-associated proteins identified various interactors typical for stress granule components and translational regulation like eukaryotic translation initiation factors or PABP proteins as well as the DNA/RNA-binding proteins ALBA1-4 (Mair *et al*., 2010; Chêne *et al*., 2012). The ALBA proteins ALBA1 and ALBA4 were previously reported to be involved in mRNA homeostasis and translational regulation, while ALBA3, which was identified by us as an interactor of MD3, has recently been reported to have nuclease activity (Vembar *et al*., 2015; Munoz *et al*., 2017; Banerjee *et al*., 2023). Noteworthy, the binding of MD3 to CITH and PUF1 was confirmed by us by co-immunoprecipitation assays. The combined protein interaction network analyses of clusters A-D let us conclude that MD3 has important roles in protein biosynthesis by interacting with both, activators and repressors of translation.

The most striking protein network cluster is cluster E, which demonstrates the interaction of MD3 with regulators of sexual development as recently identified by Russell *et al*. (2023). In this study, a genetic screen of barcode mutants was employed, which identified a set of 10 genes with roles in male and female determination in *P. berghei*. MD3 was one of three identified ZFPs and its role in sexual development was highlighted. For instance, the disruption of the *md3* gene in *P. berghei* was shown to result in a marked sex ratio shift towards females, with few fertile males, signifying its importance in male gametocytogenesis (Russell *et al*., 2023). Interestingly, we witnessed five orthologues of the 10 gametocyte development regulators of *P. berghei* in the MD3 interactome of immature P*. falciparum* gametocytes, i.e. the male development proteins MD1 and MD2, the gametocyte development protein GD1, and the female development proteins FD1 and FD2.

Best investigated in the study by Russell and colleagues (2023) is GD1, a factor for differentiation of female gametocytes. Parasites lacking GD1 show a sex ratio shift towards males, which were demonstrated to be infertile. Interaction partners of GD1, as identified by co-immunoprecipitation assays, included CCR4-NOT complex members, e.g. a NOT1-G paralogue, as well as CITH, PABP1 and PUF1, hence similar interactors as identified by us for *P. falciparum* MD3 (Russel *et al*., 2023). Interestingly, the authors claimed *P. berghei* GD1 as an orthologue of *P. falciparum* ZNF4 (see below). The two other MD3 interactors that were reported to be important for female differentiation in *P. berghei* are FD1 and FD2 and *P. berghei* parasites deficient of these proteins produce infertile female gametocytes lacking transcripts for core female markers (Russell *et al*., 2023).

Furthermore, *P. berghei* mutants deficient of MD1 and MD2 are not capable to form male gametocytes, while the females are fertile transcriptomes comparable to WT parasites (Russell *et al*., 2023). Recently, MD1 has been characterized in *P. falciparum* and was shown to be present in cytoplasmic granules, where it interacts with mRNP complexes (Gomes *et al*., 2022). The N- and C-termini of the protein define the function of MD1, as the N-terminus can solely specify a male fate, while the C-terminus comprises an OST/HTH/LOTUS domain essential for development of male gametocytes. Disruption of the *md1* gene results in impairment of male gametogenesis and also affects the morphology of microgametes. Strikingly, MD1 also shares interaction partners with MD3 that are important in translational repression as well as components of CCR4-NOT complex (Gomes *et al*., 2022). Noteworthy in this context, the MD4 orthologue in *P. falciparum*, termed, ARID, was shown to be a nuclear protein with a crucial role in male gametocyte exflagellation and macrogamete fertility (Kumar *et al*., 2022a). Finally, we confirmed the MD3 interaction with two epigenetically regulated ZFPs, i.e. RNF1, a RING-finger domain ZFP with putative E3 ubiquitin ligase activity, and the C3H1*-*ZFP ZNF4. Both ZFPs were originally identified by us in a comparative transcriptomics analysis of TSA-treated immature gametocytes, hence in the same study which enabled us to identify MD3 (Ngwa *et al*., 2017). ZNF4, which was assigned homologies to GD1 (see above), has recently been shown to have important roles in male gametocyte exflagellation by regulating male-enriched genes like ones related to microtubule/cilium morphogenesis (Hanhsen *et al*., 2022).

Noteworthy, other recent studies have also reported on RNA-binding proteins with roles in male gametogenesis. An RNA-binding protein in *P. berghei*, UIS12, was shown to be involved in gametocytogenesis with particular impact on exflagellation and hence malaria transmission (Muller *et al*., 2021). Lack of UIS12 leads to transcriptional down-regulation of various known markers of gametocytes and gametes like MiGS, P48745, P28, PPLP2 as well as various LCCL domain proteins (reviewed in Bennink *et al*., 2016; Bennink and Pradel, 2021). In addition, the transcript encoding a protein of the gametocyte inner membrane complex (IMC), IMC1j, was also affected (Muller *et al*., 2021). In relevance, the serine/arginine-rich protein kinase SRPK1, which was previously implicated in pre-mRNA splicing, was also recently assigned to male gametogenesis (Kern *et al*., 2014; Kumar *et al*., 2022b). Parasites lacking SRPK1 demonstrated the deregulation of genes coding e.g. for microtubule/cilium morphogenesis-related proteins, including IMC components.

These recent data on regulators of male gametogenesis highlight a link between RNA-binding proteins and proteins involved in IMC morphogenesis. Strikingly, the MD3 interactome of immature gametocytes as identified by us includes the PIP (Photosensitized 5-iodonaphthalene-1-azide Labelled protein-1 Interacting Proteins) members PIP1 to PIP3. assigned to the gametocyte IMC. The IMC is a flattened cisterna-like membrane compartment underneath the plasma membrane that is accompanied by microtubes and important for the falciform shape of *P. falciparum* gametocytes (Dearnley *et al*., 2012; Kono *et al*., 2013; Simon *et al*., 2013; Parkyn Schneider *et al*., 2017). During gametogenesis, the IMC is disassembled and contributes to plasma membrane restructuring of the newly formed gametes as well as to the formation of nanotubes, tubular cell-to-cell connections of gametes (Rupp *et al*., 2011; Sologub *et al*., 2011; Simon *et al*., 2013). While PIP2 and PIP3 are found in all malaria species, PIP1 is limited to human malaria parasites (Kono *et al*., 2013). With high expression of PIP1 in early gametocytes, its downregulation results in failure of *P. falciparum* gametocyte maturation as a result of impaired elongation (Parkyn Schneider *et al*., 2017).

In summary, previous studies had shown that the sex determination of *P. falciparum* gametocytes is influenced by epigenetic mechanisms during sexual commitment. However, based on current literature, the data from this study reveal that subsequent sexual development requires the transcript regulation by RNA-binding proteins. Recent research suggests that a significant network of RNA-binding proteins, including ZFPs, PUF proteins, or components of the translational control machinery, is essential for this process. During gametocyte maturation, this network of regulators appears to engage with both translational activators and repressors, thereby deciding on the fate of transcript and hence sexual development. Further studies are needed to ascertain the extent of interaction between these regulators and other protein complexes, such as the IMC proteins mentioned in this context, which also contribute to maturity and fertility of gametocytes.

## Materials and methods

### Gene Identifiers

The following PlasmoDB gene identifiers (gene IDs) are assigned to the genes and proteins investigated in this study: aldolase (PF3D7_1444800); CITH (PF3D7_1474900); falcilysin (PF3D7_1360800); histone H3 (PF3D7_0610400); MD3 (PF3D7_0315600); P25 (PF3D7_1031000); P92 (PF3D7_1364100); P230 (PF3D7_0209000); PABP1 (PF3D7_1224300); *Pf*39 (PF3D7_1108600); *Pf*AMA1 (PF3D7_1133400); *Pf*FNPA (PF3D7_1451600); PUF1 (PF3D7_0518700); RNF1 (PF3D7_0314700); ZNF4 (PF3D7_1134600).

### Antibodies

In this study, the following antibodies were used: rabbit polyclonal antisera against HA (Sigma-Aldrich), P230 (Biogenes), P25 (ATCC, Manassas, USA), P92 (Musabyimana *et al*., 2022), *Pf*39 (Davids Biotechnology, Regensburg, Germany), *Py*PABP1 (Minns *et al*., 2018), and *Py*CITH (Bennink *et al*., 2018); mouse polyclonal antisera against falcilysin (Weißbach *et al*., 2017); rabbit anti-H3K4me3 antibody (abcam, Cambridge, UK), monoclonal rat anti-HA antibody (Roche, Basel, Switzerland) and mouse anti-GFP antibody (Roche, Basel, Switzerland). Mouse antisera against MD3 and RNF1 were generated as described below. The following dilutions were used for IFAs: rabbit anti-P230 (1:200), rabbit anti-P92 (1:200), mouse anti-MD3 (1:20), rabbit anti-P25 (1:200), rabbit anti-*Pf*39 (1:200) mouse anti-GFP (1:200), rat anti-HA (1:50). For WB analysis, the following dilutions were used: rat anti-HA (1:500), rabbit anti-HA (1:1000), rabbit anti-*Pf*39 (1:10,000), mouse anti-GFP (1:1,000), mouse anti-MD3 (1:300), mouse anti-RNF1 (1:1,000), rabbit anti-CITH (1:1,000), rabbit anti-PABP1 (1:1,000).

### Bioinformatics

The 3D structure of MD3 was predicted using the Alphafold programme (https://alphafold.ebi.ac.uk/; Jumper *et al*., 2021; Varadi *et al*., 2022). Predictions of gene expression and protein properties and function were made using the database PlasmoDB (http://plasmoDB.org; Aurrecoechea *et al.,* 2009); the peak transcript expression of candidate genes was analyzed using table “Transcriptomes of 7 sexual and asexual life stages” (López-Barragán *et al.,* 2011) and sex specificity was predicted using table “Gametocyte Transcriptomes” (Lasonder *et al.,* 2016) of the PlasmoDB database. The gene ontology enrichment (GO) analysis was performed using the ShinyGO 0.77 (Ge *et al*., 2020) with a p-value cut-off of 0.05. A network analysis was conducted using the STRING database (version 11.0) (Szklarczyk *et al.,* 2019), using default settings, including text mining options and a confidence of 0.4.

### Parasite culture

The WT NF54 and all generated mutant parasite lines were cultivated *in vitro* in human blood group A+ erythrocytes as previously described (Ifediba & Vanderberg 1981). Asexual blood stages and gametocytes were maintained in RPMI 1640/HEPES medium (Gibco; Thermo Fisher Scientific; Waltham, US) supplemented with 10% (v/v) heat inactivated human A+ serum, 50 μg/ml hypoxanthine (Sigma-Aldrich; Taufkirchen, DE) and 10 μg/ml gentamicin (Gibco; Thermo Fisher Scientific; Waltham, US). For cultivation of the mutant parasite lines, the selection drug WR99210 (Jacobus Pharmaceutical Company; Princeton, US) was added at a final concentration of 4.0 nM. All cultures were kept at 37°C in an atmosphere of 5% O2 and 5% CO2 in N2. Synchronization of cultures was carried out by repeated 5% sorbitol treatment as described (Lambros and Vanderberg, 1979). For MD3 knockdown, MD3-HA-*glmS* cultures were incubated with a complete medium supplemented with 5 mM glucosamine hydrochloride (GlcN; D-(+)-glucosamine hydrochloride; Sigma-Aldrich, Taufkirchen, DE). For generation of gametocytes, the cultures with mainly ring stages at high parasitemia were used and gametocytogenesis was induced by the addition of lysed RBCs. After 24 h, lysed RBCs were washed and the culture medium was supplemented with 50 mM N-acetyl glucosamine (GlcNac; Carl Roth, Karlsruhe, Germany) for ∼5 days to kill the asexual blood stages (Fivelman *et al*., 2007). For knockdown assays, heparin sulfate (Sigma-Aldrich, Taufkirchen, Germany), with final concentration of 20 U/ml was used to kill asexual blood stages as described (Miao *et al*., 2013b). Gametocytes were enriched via Percoll (Cytiva; Washington DC, US) gradient centrifugation as described previously (Kariuki *et al.,* 1998). Gametogenesis was induced by adding xanthurenic acid in a final concentration of 100 μM dissolved in 1% (v/v) 0.5 M NH4OH/ddH2O and incubation for 15 or 30 min at room temperature (RT). For studies on epigenetic regulation, Percoll-enriched immature gametocytes were incubated with 0.26 μM TSA (Cell Signaling Technology, Danvers, US) in 0.5% (v/v) ethanol for 24 h at at 37°C. Erythrocyte concentrate and serum from humans were procured from the Department of Transfusion Medicine (University Hospital Aachen, Germany). All work with human blood was approved by the University Hospital Aachen Ethics commission, the donors remained anonymous and serum samples were pooled.

### Generation of mouse antisera directed against MD3 and RNF1

Recombinant peptides, corresponding to portions of MD3 (aa 31-359; Fig. 1A) and RNF1 (aa 294-548) were expressed in the *Escherichia coli* system, fused to a maltose-binding tag using the pMAL™c5X-vector (New England Biolabs, Ipswich, US). Briefly, the coding sequences were amplified using gene-specific primers (for primer sequences, see Table S5) and the recombinant proteins were expressed in *E. coli* BL21 (DE3) RIL cells following the manufacturer’s protocol (Invitrogen, Karlsruhe, DE). Each protein was then isolated and affinity-purified using an amylose resin according to the manufactureŕs protocol (New England Biolabs, Ipswich, US) and the concentration was determined using the Bradford technique. Six weeks old female NMRI mice (Charles River Laboratories, Wilmington, US) were subcutaneously injected with 100 µg of the pure recombinant protein emulsified in Freund’s incomplete adjuvant (Sigma Aldrich, Taufkirchen, DE) followed by a boost after 4 weeks. At day 10 after the boost, immune sera were collected via heart puncture after anesthetizing the mice by intraperitoneal injection of a mixture of ketamine and xylazine according to the manufacturer’s protocol (Sigma Aldrich, Taufkirchen, DE). The immune sera of three immunized mice were pooled; sera of three non-immunized mice were used as negative control. Experiments in mice were approved by the animal welfare committee of the District Council of Cologne, Germany (ref. no. 84-02.05.30.12.097 TVA).

### Generation of the MD3-HA-*glmS* and PUF1-HA-*glmS* parasite lines

The MD3-HA-*glmS* and PUF1-HA-*glmS* transgenic parasite lines were generated via single-crossover homologous recombination, using vector pSLI-HA-*glmS* (Fig. S3A; Musabyimana *et al.,* 2022). In this method, we modified the plasmid to contain a homology block from the 3’ end of the respective genes without the stop codon (for primer sequence, see Table S5), thereby enabling the coding region for genes to be fused at the 3’-region to a HA-encoding sequence followed by the *glmS*-ribozyme sequence in the vector. Ligation of the insert and vector backbone was mediated by NotI and XmaI restriction sites. A WT NF54 culture with mainly ring stages was used to transfect with 100 µg plasmid DNA in transfection buffer via electroporation (310 V, 950 μF, 12 ms; Bio-Rad gene-pulser Xcell) as described earlier (e.g. Wirth *et al.,* 2014; Ngwa *et al.,* 2017). A mock control lacking plasmid DNA was electroporated using transfection buffer and was cultured in medium with and without WR99210. After 6 h of transfection, the integrated parasites were selected using WR99210 at a final concentration of 4 nM. WR99210-resistant parasites appeared at ∼21 days post-transfection and the parasites were treated with medium supplemented with 550 μg/ml neomycin (G418 disulfate salt; Sigma-Aldrich; Taufkirchen, DE) for two weeks to remove WT NF54 parasites from the culture. The gDNA was isolated from the transgenic parasite lines using the NucleoSpin Blood Kit (Macherey-Nagel; Dueren, DE) following the manufacturer’s protocol and used as template to verify the successful vector integration by diagnostic PCR (for primer sequence, see Table S5).

### Generation of the MD3-GFP-BirA parasite lines

The MD3-GFP-BirA parasite lines were generated, using vectors pARL-*pfama1*/*pffnpa*-GFP-BirA (Musabyimana *et al.,* 2022). The sequence for full-length *md3* was cloned into the vector pARL-GFP-BirA (for primer sequences, see Table S5) under the control of the *pfama1* and *pffnpa* promoters, using vectors pARL-*pfama1*-GFP-BirA and pARL-*pffnpa*-GFP-BirA, as described (Fig. S9A; Musabyimana *et al*., 2022). The restriction enzymes KpnI and AvrII were used to ligate the insert and the vector backbone. The constructed plasmids were then used to transfect WT NF54 parasites as described above. For selection of parasites episomally overexpressing the proteins, WR99210 was added to a final concentration of 4 nM and successful uptake of the vector was confirmed by diagnostic PCR (for primer sequences, see Table S5).

### Indirect immunofluorescence assay

Parasite cultures containing mixed asexual blood stages and gametocytes of WT NF54 as well as lines MD3-HA-*glmS*, MD3-*pfama1*-GFP-BirA, and MD3-*pffnpa*-GFP-BirA were coated on glass slides as cell monolayers and then air-dried. After fixation in methanol at −80 °C for 10 min, the cells were serially incubated in 0.01% (w/v) saponin/0.5% (w/v) BSA/PBS and 1% (v/v) neutral goat serum (Sigma-Aldrich)/PBS for 30 min at RT to facilitate membrane permeabilization and blocking of non-specific binding. The primary antibodies were diluted in 3% (w/v) BSA/PBS and were added to the slide for 2 h incubation at 37°C. The slides were washed 3x with PBS and incubated with the secondary antibody for 1 h at 37°C. Following 2x washing with PBS, the incubation with the second primary antibody and the corresponding visualization with the second secondary antibody were carried out as described above. The nuclei were stained with Hoechst 33342 staining solution for 2 min at RT (1:5,000 in 1x PBS). The cells were mounted with anti-fading solution (Citifluor Limited; London, UK), covered with a coverslip and sealed airtight with nail polish. The parasites were visualized by conventional fluorescence microscopy using a Leica DM5500 B (Leica; Wetzlar, DE) microscope. The following secondary antibodies were used: Anti-mouse Alexa Fluor 488, anti-rabbit Alexa Fluor 488, anti-rat Alexa Fluor 488, anti-mouse Alexa Fluor 594, anti-rabbit Alexa Fluor 594, (1:1,000; all fluorophores from Invitrogen Molecular Probes; Eugene, US, or Sigma-Aldrich; Taufkirchen, DE); further Alexa Fluor 594 streptavidin (1:500; Invitrogen Molecular Probes; Eugene, US) was used. Alternatively, the asexual blood stages were stained with 0.01% (w/v) Evans Blue (Sigma-Aldrich; Taufkirchen, DE)/PBS for 3 min at RT followed by 5 min washing with PBS. Images were processed using the Adobe Photoshop CS software.

### Subcellular fractionation

Percoll-purified immature gametocytes of WT NF54 and the MD3-HA-*glmS* line were lysed by using lysis buffer (20 mM HEPES (pH 7.8), 10 mM KCl, 1 mM EDTA, 1 mM dithiothreitol (DTT), 1 mM PMSF, 1% (v/v) Triton X-100), with addition of protease inhibitor cocktail (complete EDTA-free, Roche) and incubated for 10 min on ice. The lysates were carefully centrifuged at 2,500 × g for 5 min at 4 °C, the cytosolic proteins were harvested with the supernatant and stored at −80 °C. Following several washing steps for the remaining pellet with the lysis buffer, nuclear proteins were extracted with about 2 volumes of the extraction buffer (20 mM HEPES (pH 7.8), 800 mM KCl, 1 mM EDTA, 1 mM DTT, and 1 mM PMSF), after addition of the protease inhibitor cocktail. Following incubation for 30 min, while rotating at 4°C, the extract was cleared by centrifugation at 13,000 xg for 30 min at 4°C. Finally, the supernatants were diluted in a 1:1 ratio with dilution buffer (20 mM HEPES (pH 7.8); 1 mM EDTA; 1 mM DTT; 30% (v/v) glycerol) and stored at −80 °C. The individual fractions were subjected to WB as described below.

### Co-immunoprecipitation assay

Percoll-purified immature gametocytes of WT NF54 and the transgenic lines MD3-HA-*glmS*, ZNF4-HA-*glmS* (Hanhsen *et al*., 2022) and PUF1-HA-*glmS* were lysed with RIPA buffer (150 mM NaCl, 1% (v/v) Triton X-100, 0.5% (w/v) sodium deoxycholate, 0.1% (w/v) SDS, 50 mM Tris buffer). Following incubation on ice for 15 min, three sessions of sonication were applied to each sample (30sec/ 50% and 0.5 cycles). After centrifugation (16,000 x g for 10 min at 4 °C), the supernatant was incubated with 5% (v/v) pre-immune rabbit or rat sera for 30 min, followed by incubation with 20 μl of protein G-beads (Roche) for 1 h on a rotator at 4°C. After centrifugation (3,500 x g for 5 min at 4°C), the supernatant was incubated for 1 h at 4°C with 5% (v/v) monoclonal rat or polyclonal rabbit anti-HA antisera. Afterwards, a volume of 30 μl protein G-beads was added and kept on rotation overnight at 4°C. Following centrifugation (3,500 x g for 5 min at 4 °C), beads were first washed with ice cold RIPA buffer and then with PBS for five times. The beads were resuspended in an equal volume of loading buffer and the samples were subjected to WB analysis as described below.

### Western blotting

Asexual blood stage parasites of WT NF54 and lines MD3-HA-*glmS*, MD3-*pfama1*-GFP-BirA and PUF1-HA-*glmS* were harvested from mixed or synchronized cultures, while gametocytes of WT NF54, or lines MD3-HA-*glmS*, MD3-*pffnpa*-GFP-BirA and PUF1-HA-*glmS* were enriched by Percoll purification. The infected RBCs were lysed with 0.05% (w/v) saponin/PBS for 10 min at 4°C to release the parasites, then washed with PBS, and resuspended in lysis buffer (0.5% (v/v) Triton X-100, 4% (w/v) SDS in PBS) supplemented with protease inhibitor cocktail. After adding 5x SDS-PAGE loading buffer containing 25 mM DTT, the lysates were heat-denatured for 10 min at 95°C. Lysates were separated via SDS-PAGE and transferred to Hybond ECL nitrocellulose membranes (Amersham Biosciences, Buckinhamshire, UK) following the manufacturer’s protocol. Non-specific binding sites were blocked by incubation with 5% (w/v) skim milk and 1% (w/v) BSA in Tris buffer (pH 7.5) for 1 h at 4°C. For immunodetection, membranes were incubated overnight at 4°C with the primary antibody in 3% (w/v) skim milk/TBS. The membranes were washed 3x each with 3% (w/v) skim milk/TBS and 3% (w/v) skim milk/0.1 % (v/v) Tween/TBS and then incubated for 1 h at RT with goat anti-mouse, anti-rabbit or anti-rat alkaline phosphatase-conjugated secondary antibodies (dilution 1:10,000, Sigma-Aldrich; Taufkirchen, DE) in 3% (w/v) skim milk/TBS. The membranes were developed in a NBT/BCIP solution (nitroblue tetrazolium chloride/5-bromo-4-chloro-3-indoxyl phosphate; Roche; Basel, CH) for up to 20 min at RT. For the detection of biotinylated proteins, the blocking step was performed overnight at 4°C in 5% (w/v) skim milk/TBS and the membrane was washed 5x with 1x TBS before incubation with streptavidin-conjugated alkaline phosphatase (dilution 1:1,000, Sigma-Aldrich; Taufkirchen, DE) in 5% (w/v) BSA/TBS for 1 h at RT. Blots were scanned and processed using the Adobe Photoshop CS software. Band intensities were measured using the ImageJ program version 1.51f.

### Asexual blood stage replication assay

Asexual blood stage cultures of line MD3-HA-*glmS* or WT NF54 were tightly synchronized and set to an initial parasitemia of 0.25% ring stages. The cultures were then continuously treated with 5 mM GlcN for transcript knockdown. Untreated cultures were used for control. Giemsa-stained thin blood smears were prepared every 24 h over a time period of 96 h at five different time points (0, 24, 48, 72 and 96 h post-seeding). At each time point, the parasitemia was determined microscopically at 1,000-fold magnification by counting the percentage of parasites in 1,000 RBCs. Blood stages (ring, trophozoites, schizonts) present in the cultures were identified at each time point by counting 50 infected RBC (iRBCs) per setting. For each assay, three experiments were performed, each in triplicate. Data analysis was performed using MS Excel 2016 and GraphPad Prism 5.

### Gametocyte development assay

Tightly synchronized ring stage parasite cultures of line MD3-HA-*glmS* and WT NF54 were induced at a parasitemia of 7.5% with lysed RBCs for 24 h (Schneweis *et al*., 1991; Ngwa *et al*., 2017). The cultures were subsequently maintained in cell culture medium supplemented with 20 U/ml heparin to kill the asexual blood stages for 4 days. The cultures were then treated with 5mM GlcN for transcript knockdown until day 13. Untreated cultures were used as control. Giemsa-stained thin blood smears were prepared at days 3, 5, 7, 9, 11 and 13 post-induction. Gametocytemia was determined per 1,000 RBCs and the numbers of gametocyte stages II to V were quantified microscopically for 50 iRBCs per setting. For each assay, three experiments were performed, each in triplicate. Data analysis was performed using MS Excel 2016 and GraphPad Prism 5.

### Exflagellation assay

Tightly synchronized ring stage parasite cultures of line MD3-HA-*glmS* and WT NF54 were induced for gametocytogenesis by addition of lysed blood as described above. Cells were washed and treated with 20 U/ml heparin for two days. On day 7 post-induction, 5 mM GlcN was added for 4 days for transcript knockdown. Untreated cultures were used for control. The cultures were allowed to recover for a period of 2 days prior to the exflagellation assays. On day 13, exflagellation of male gametocytes was investigated, using light microscopy. For this, all four cultures were adjusted to total RBC numbers and the number of mature gametocytes were counted for each culture via Giemsa-stained smears. For gametocyte activation, 100 µl of each gametocyte culture was activated by addition of 100 µM XA for 15 min at RT. The number of exflagellation centers were counted at 400-fold magnification in 30 optical fields in triplicate using a microscope and the results were adjusted to the number of mature gametocytes determined before gametocyte activation. Data analysis was done by using GraphPad Prism 5.

### Preparation of samples for BioID analysis

Highly synchronized ring stage parasite cultures of line MD3-*pfama1*-GFP-BirA and Percoll-purified immature gametocyte cultures of the MD3-*pffnpa*-GFP-BirA parasite line as well as WT NF54 as control were treated with biotin (Sigma-Aldrich; Taufkirchen, DE), at a final concentration of 50 µM for 24 h to induce the biotinylation of proximal proteins by the BirA ligase. After treatment, the RBCs were lysed with 0.05% (w/v) saponin and the parasites were resuspended in 100 µl binding buffer (Tris buffer containing 1% (v/v) Triton X-100 and protease inhibitor). The sample was sonicated on ice (2 x 60 pulses at 30% duty cycle) and another 100 µl of ice-cold binding buffer was added. After a second session of sonification, cell debris was pelleted by centrifugation (5 min, 16,000x *g*, 4°C). The supernatant was mixed with pre-equilibrated Cytiva Streptavidin Mag Sepharose Magnet-Beads (Cytiva; Washington DC, US) in a low-binding reaction tube. Incubation was performed with slow end-over-end mixing over night at 4°C. The beads were washed 6x with 500 μl washing buffer (3x: RIPA buffer containing 0.03% (w/v) SDS, followed by 3x 25 mM Tris buffer (pH 7.5)) and were resuspended in 45 μl elution buffer (1% (w/v) SDS/5 mM biotin in Tris buffer (pH 7.5)), followed by an incubation for 5 min at 95°C. The supernatant was transferred into a new reaction tube and stored at −20°C. For each culture, three independent samples were collected.

### Proteolytic digestion

Samples were processed by single-pot solid-phase-enhanced sample preparation (SP3) as described before (Hughes *et al.,* 2014; Sielaff *et al.,* 2017). In brief, proteins bound to the streptavidin beads were released by incubating the samples for 5 min at 95° in an SDS-containing buffer (1% (w/v) SDS, 5 mM biotin in water/Tris buffer, pH 8.0). After elution, proteins were reduced and alkylated, using DTT and iodoacetamide (IAA), respectively. Afterwards, 2 µl of carboxylate-modified paramagnetic beads (Sera-Mag Speed Beads, GE Healthcare; Chicago, US; 0.5 μg solids/μl in water as described by Hughes *et al.,* 2014) were added to the samples. After adding acetonitrile to a final concentration of 70% (v/v), samples were allowed to settle at RT for 20 min. Subsequently, beads were washed twice with 70% (v/v) ethanol in water and once with acetonitrile. The beads were resuspended in 50 mM NH4HCO3 supplemented with trypsin (Mass Spectrometry Grade, Promega; Madison, US) at an enzyme-to-protein ratio of 1:25 (w/w) and incubated overnight at 37°C. After overnight digestion, acetonitrile was added to the samples to reach a final concentration of 95% (v/v) followed by incubation at RT for 20 min. To increase the yield, supernatants derived from this initial peptide-binding step were additionally subjected to the SP3 peptide purification procedure (Sielaff *et al.,* 2017). Each sample was washed with acetonitrile. To recover bound peptides, paramagnetic beads from the original sample and corresponding supernatants were pooled in 2% (v/v) dimethyl sulfoxide (DMSO) in water and sonicated for 1 min. After 2 min of centrifugation at 14,000xg and 4°C, supernatants containing tryptic peptides were transferred into a glass vial for MS analysis and acidified with 0.1% (v/v) formic acid.

### Liquid chromatography-mass spectrometry (LC-MS) analysis

Tryptic peptides were separated using an Ultimate 3000 RSLCnano LC system (Thermo Fisher Scientific; Waltham, US) equipped with a PEPMAP100 C18 5 µm 0.3 x 5 mm trap (Thermo Fisher Scientific; Waltham, US) and an HSS-T3 C18 1.8 μm, 75 μm x 250 mm analytical reversed-phase column (Waters Corporation; Milford, US). Mobile phase A was water containing 0.1% (v/v) formic acid and 3% (v/v) DMSO. Peptides were separated by running a gradient of 2–35% mobile phase B (0.1% (v/v) formic acid, 3% (v/v) DMSO in ACN) over 40 min at a flow rate of 300 nl/min. Total analysis time was 60 min including wash and column re-equilibration steps. Column temperature was set to 55°C. Mass spectrometric analysis of eluting peptides was conducted on an Orbitrap Exploris 480 (Thermo Fisher Scientific; Waltham, US) instrument platform. Spray voltage was set to 1.8 kV, the funnel RF level to 40, and heated capillary temperature was at 275°C. Data were acquired in data-dependent acquisition (DDA) mode targeting the 10 most abundant peptides for fragmentation (Top10). Full MS resolution was set to 120,000 at *m/z* 200 and full MS automated gain control (AGC) target to 300% with a maximum injection time of 50 ms. Mass range was set to *m/z* 350–1,500. For MS2 scans, the collection of isolated peptide precursors was limited by an ion target of 1 × 10^5^ (AGC tar-get value of 100%) and maximum injection times of 25 ms. Fragment ion spectra were acquired at a resolution of 15,000 at *m/z* 200. Intensity threshold was kept at 1E4. Isolation window width of the quadrupole was set to 1.6 *m/z* and normalized collision energy was fixed at 30%. All data were acquired in profile mode using positive polarity. Each sample was analyzed in three technical replicates.

### Data analysis and label-free quantification

DDA raw data acquired with the Exploris 480 were processed with MaxQuant (version 2.0.1; Cox & Mann 2008; Cox *et al.,* 2014), using the standard settings and label-free quantification (LFQ) enabled for each parameter group, i.e. control and affinity-purified samples (LFQ min ratio count 2, stabilize large LFQ ratios disabled, match-between-runs). Data were searched against the forward and reverse sequences of the *P. falciparum* proteome (UniProtKB/TrEMBL, 5,445 entries, UP000001450, released April 2020) and a list of common contaminants. For peptide identification, trypsin was set as protease allowing two missed cleavages. Carbamidomethylation was set as fixed and oxidation of methionine as well as acetylation of protein N-termini as variable modifications. Only peptides with a minimum length of 7 amino acids were considered. Peptide and protein false discovery rates (FDR) were set to 1%. In addition, proteins had to be identified by at least two peptides. Statistical analysis of the data was conducted using Student’s t-test, which was corrected by the Benjamini-Hochberg (BH) method for multiple hypothesis testing (FDR of 0.01). In addition, proteins in the affinity-enriched samples had to be identified in all three biological replicates and show at least a two-fold enrichment compared to the controls. The datasets of protein hits were further edited by verification of the gene IDs and gene names via the PlasmoDB database (www.plasmodb.org; Aurrecoechea *et al.,* 2009). PlasmoDB gene IDs were extracted from the fasta headers provided by mass spectrometry and verified manually. Once an initial list of significantly enriched proteins had been established, proteins with a putative signal peptide were excluded.

### Data availability

The mass spectrometry proteomics data have been deposited to the ProteomeXchange Consortium (http://proteomecentral.proteomexchange.org) via the jPOST partner repository (Vizcaino *et al*., 2013) with the dataset identifiers PXD042976 (ProteomeXchange) and JPST002199 (jPOST).

### Statistical Analysis

Data are presented as mean ± SD. Statistical differences were determined using one-way ANOVA with post hoc Bonferroni multiple comparison test or unpaired two-tailed Student’s *t*-test, as indicated. The *p*-values < 0.05 were considered statistically significant. Significances were calculated using GraphPad Prism 5 and are represented in the figures as follows: * *p* < 0.05; ** *p* < 0.01; *** *p* < 0.001.

## Acknowledgements

The authors acknowledge funding by the Deutsche Forschungsgemeinschaft (DFG; project grants PR905/15-1 and PR905/20-1 to GP and NG170/1-1 to CJN; DFG priority programme SPP 2225 grants PR905/19-1 to GP, and TE599/9-1 to ST). AF and JPM were supported by stipends from the Deutscher Akademischer Austauschdienst (DAAD).

## References

1. Aurrecoechea, C., Brestelli, J., Brunk, B. P., Dommer, J., Fischer, S., Gajria, B., … Wang, H. (2009). PlasmoDB: a functional genomic database for malaria parasites. Nucleic Acids Research, 37, D539–D543. https://doi.org/10.1093/nar/gkn814

2. Balbin, J. M., Heinemann, G. K., Yeoh, L. M., Gilberger, T. W., Armstrong, M., Duffy, M. F., … Wilson, D. W. (2023). Characterisation of PfCZIF1 and PfCZIF2 in Plasmodium falciparum asexual stages. International Journal for Parasitology, 53, 27–41. https://doi.org/10.1016/j.ijpara.2022.09.008

3. Bancells, C., Llora-Batlle, O., Poran, A., Notzel, C., Rovira-Graells, N., Elemento, O., … Cortes, A. (2019). Revisiting the initial steps of sexual development in the malaria parasite Plasmodium falciparum. Nature Microbiolology, 4, 144–154. https://doi.org/10.1038/s41564-018-0291-7

4. Banerjee, C., Nag, S., Goyal, M., Saha, D., Siddiqui, A. A., Mazumder, S., Bandyopadhyay, U. (2023). Nuclease activity of Plasmodium falciparum Alba family protein PfAlba3. Cell Reports, 42, 112292. https://doi.org/10.1016/j.celrep.2023.112292

5. Baumgarten, S., Bryant, J. M., Sinha, A., Reyser, T., Preiser, P. R., Dedon, P. C., & Scherf, A. (2019). Transcriptome-wide dynamics of extensive m(6)A mRNA methylation during Plasmodium falciparum blood-stage development. Nature Microbiology, 4, 2246–2259. https://doi.org/10.1038/s41564-019-0521-7

6. Bennink, S., Kiesow, M. J., & Pradel, G. (2016). The development of malaria parasites in the mosquito midgut. Cellular Microbiology, 18, 905–918. https://doi.org/10.1111/cmi.12604

7. Bennink, S., & Pradel, G. (2019) The molecular machinery of translational control in malaria parasites. Molecular Microbiology, 112, 111372. https://doi.org/10.1111/mmi.14388

8. Bennink, S., & Pradel, G. (2021) Vesicle dynamics during the egress of malaria gametocytes from the red blood cell. Molecular and Biochemical Parasitology, 234, 1658–1673. https://doi.org/10.1016/j.molbiopara.2021.111372.

9. Bennink, S., von Bohl, A., Ngwa, C. J., Henschel, L., Kuehn, A., Pilch, N., … Pradel, G. (2018). A seven-helix protein constitutes stress granules crucial for regulating translation during human-to-mosquito transmission of Plasmodium falciparum. PLoS Pathogens, 14, e1007249. https://doi.org/10.1371/journal.ppat.1007249

10. Birnbaum, J., Flemming, S., Reichard, N., Soares, A. B., Mesen-Ramirez, P., Jonscher, E., … Spielmann, T. (2017). A genetic system to study Plasmodium falciparum protein function. Nature Methods, 14, 450–456. https://doi.org/10.1038/nmeth.4223

11. Brancucci, N. M. B., Bertschi, N. L., Zhu, L., Niederwieser, I., Chin, W. H., Wampfler, R., … Voss, T.S. (2014). Heterochromatin protein 1 secures survival and transmission of malaria parasites. Cell Host and Microbe, 16, 165–176. https://doi.org/10.1016/j.chom.2014.07.004

12. Bunnik, E. M., Batugedara, G., Saraf, A., Prudhomme, J., Florens, L., & Le Roch, K. G. (2016). The mRNA-bound proteome of the human malaria parasite Plasmodium falciparum. Genome Biology, 17, 147. https://doi.org/10.1186/s13059-016-1014-0

13. Chene, A., Vembar, S. S., Riviere, L., Lopez-Rubio, J. J., Claes, A., Siegel, T. N., Scherf, A. (2012). PfAlbas constitute a new eukaryotic DNA/RNA-binding protein family in malaria parasites. Nucleic Acids Research, 40, 3066–3077. https://doi.org/10.1093/nar/gkr1215

14. Coleman, B. I., Skillman, K. M., Jiang, R. H. Y., Childs, L. M., Altenhofen, L. M., Ganter, M., … Duraisingh, M. T. (2014). A Plasmodium falciparum histone deacetylase regulates antigenic variation and gametocyte conversion. Cell Host and Microbe, 16, 177–186. https://doi.org/10.1016/j.chom.2014.06.014

15. Cox, J., Hein, M. Y., Luber, C. A., Paron, I., Nagaraj, N., & Mann, M. (2014). Accurate proteome-wide label-free quantification by delayed normalization and maximal peptide ratio extraction, termed MaxLFQ. Mol Cell Proteomics 13, 2513–2526. https://doi.org/10.1074/mcp.M113.031591

16. Cox, J., & Mann, M. (2008). MaxQuant enables high peptide identification rates, individualized p.p.b.-range mass accuracies and proteome-wide protein quantification. Nature Biotechnology, 26, 1367–1372. https://doi.org/10.1038/nbt.1511

17. Cui, L., Mharakurwa, S., Ndiaye, D., Rathod, P. K., & Rosenthal, P. J. (2015). Antimalarial Drug Resistance: Literature Review and Activities and Findings of the ICEMR Network. American Journal of Tropical Medicine and Hygiene, 93, 57–68. https://doi.org/10.4269/ajtmh.15-0007

18. Dearnley, M. K., Yeoman, J. A., Hanssen, E., Kenny, S., Turnbull, L., Whitchurch, C. B., … Dixon, M. W. (2012). Origin, composition, organization and function of the inner membrane complex of Plasmodium falciparum gametocytes. Journnal of Cell Science, 125, 2053–2063. https://doi.org/10.1242/jcs.099002

19. Filarsky, M., Fraschka, S. A., Niederwieser, I., Brancucci, N. M. B., Carrington, E., Carrio, E.,… Voss, T. S. (2018). GDV1 induces sexual commitment of malaria parasites by antagonizing HP1-dependent gene silencing. Science, 359, 1259–1263. https://doi.org/10.1126/science.aan6042

20. Fivelman, Q. L., McRobert, L., Sharp, S., Taylor, C. J., Saeed, M., Swales, C. A., … Baker, D.A. (2007). Improved synchronous production of Plasmodium falciparum gametocytes in vitro. Molecular and Biochemical Parasitology, 154, 119–123. https://doi.org/10.1016/j.molbiopara.2007.04.008

21. Furuya, T., Mu, J., Hayton, K., Liu, A., Duan, J., Nkrumah, L., … Su, X. Z. (2005). Disruption of a Plasmodium falciparum gene linked to male sexual development causes early arrest in gametocytogenesis. Proceedings of the National Academy of Sciences USA, 102, 16813–16818. https://doi.org/10.1073/pnas.050185810

22. Ge, S. X., Jung, D., & Yao, R. (2020). ShinyGO: a graphical gene-set enrichment tool for animals and plants. Bioinformatics, 36, 2628–2629. https://doi.org/10.1093/bioinformatics/btz931

23. Gomes, A. R., Marin-Menendez, A., Adjalley, S. H., Bardy, C., Cassan, C., Lee, M. C. S., & Talman, A. M. (2022). A transcriptional switch controls sex determination in Plasmodium falciparum. Nature, 612, 528–533. https://doi.org/10.1038/s41586-022-05509-z

24. Goyal, M., Simantov, K., & Dzikowski, R. (2022). Beyond splicing: serine-arginine proteins as emerging multifaceted regulators of RNA metabolism in malaria parasites. Current Opinion in Microbiology, 70, 102201. https://doi.org/10.1016/j.mib.2022.102201

25. Hanhsen, B., Farrukh, A., Pradel, G., & Ngwa, C. J. (2022). The Plasmodium falciparum CCCH Zinc Finger Protein ZNF4 Plays an Important Role in Gametocyte Exflagellation through the Regulation of Male Enriched Transcripts. Cells, 11, 1666. https://doi.org/10.3390/cells11101666

26. Hart, K. J., Oberstaller, J., Walker, M. P., Minns, A. M., Kennedy, M. F., Padykula, I., … Lindner, S. E. (2019). Plasmodium male gametocyte development and transmission are critically regulated by the two putative deadenylases of the CAF1/CCR4/NOT complex. PLoS Pathogens, 15. e1007164. https://doi.org/10.1371/journal.ppat.1007164

27. Hart, K. J., Power, B. J., Rios, K. T., Sebastian, A., & Lindner, S. E. (2021). The Plasmodium NOT1-G paralogue is an essential regulator of sexual stage maturation and parasite transmission. PLoS Biology, 19, e3001434. https://doi.org/10.1371/journal.pbio.3001434

28. Hughes, C. S., Foehr, S., Garfield, D. A., Furlong, E. E., Steinmetz, L. M., & Krijgsveld, J. (2014). Ultrasensitive proteome analysis using paramagnetic bead technology. Molecular Systems Biology, 10, 757. https://doi.org/10.15252/msb.20145625

29. Ifediba, T., & Vanderberg, J. P. (1981). Complete in vitro maturation of Plasmodium falciparum gametocytes. Nature, 294, 364–366. https://doi.org/10.1038/294364a0

86. Josling, G. A., Russell, T. J., Venezia, J., Orchard, L., van Biljon, R., Painter, H. J., & Llinas, M. (2020). Dissecting the role of PfAP2-G in malaria gametocytogenesis. Nature Communications, 11, 1503. https://doi.org/10.1038/s41467-020-15026-0

30. Jumper, J., Evans, R., Pritzel, A., Green, T., Figurnov, M., Ronneberger, O., … Hassabis, D. (2021). Highly accurate protein structure prediction with AlphaFold. Nature, 596, 583–589. https://doi.org/10.1038/s41586-021-03819-2

31. Kafsack, B. F., Rovira-Graells, N., Clark, T. G., Bancells, C., Crowley, V. M., Campino, S. G., … Llinas, M. (2014). A transcriptional switch underlies commitment to sexual development in malaria parasites. Nature, 507, 248–252. https://doi.org/10.1038/nature12920

32. Kariuki, M.M., Kiaira, J.K., Mulaa, F.K., Mwangi, J.K., Wasunna, M.K., & Martin, S.K. (1998). Plasmodium falciparum: purification of the various gametocyte developmental stages from in vitro-cultivated parasites. The American journal of tropical medicine and hygiene, 4, 505–508. https://doi.org/10.4269/ajtmh.1998.59.505

33. Kern, S., Agarwal, S., Huber, K., Gehring, A. P., Strodke, B., Wirth, C. C., Pradel, G. (2014). Inhibition of the SR protein-phosphorylating CLK kinases of Plasmodium falciparum impairs blood stage replication and malaria transmission. PLoS One, 9, e105732. https://doi.org/10.1371/journal.pone.0105732

34. Kono, M., Herrmann, S., Loughran, N. B., Cabrera, A., Engelberg, K., Lehmann, C., … Gilberger, T. W. (2012). Evolution and architecture of the inner membrane complex in asexual and sexual stages of the malaria parasite. Molecular Biology and Evolution, 29, 2113–2132. https://doi.org/10.1093/molbev/mss081

35. Kumar, S., Baranwal, V. K., Haile, M. T., Oualim, K. M. Z., Abatiyow, B. A., Kennedy, S. Y., … Kappe, S. H. I. (2022a). PfARID Regulates P. falciparum Malaria Parasite Male Gametogenesis and Female Fertility and Is Critical for Parasite Transmission to the Mosquito Vector. mBio, 13, e0057822. https://doi.org/10.1128/mbio.00578-22

36. Kumar, S., Baranwal, V. K., Leeb, A. S., Haile, M. T., Oualim, K. M. Z., Hertoghs, N., … Kappe, S. H.I. (2022b). PfSRPK1 Regulates Asexual Blood Stage Schizogony and Is Essential for Male Gamete Formation. Microbiology Spectrum, 10, e0214122. https://doi.org/10.1128/spectrum.02141-22

37. Lal, K., Delves, M.J., Bromley, E., Wastling, J.M., Tomley, F.M., Sinden, R.E. (2009) Plasmodium male development gene-1 (mdv-1) is important for female sexual development and identifies a polarised plasma membrane during zygote development. International Journal for Parasitology, 39, 755–761. https://doi.org/10.1016/j.ijpara.2008.11.008

38. Lambros, C., & Vanderberg, J. P. (1979). Synchronization of Plasmodium falciparum erythrocytic stages in culture. Journal of Parasitology, 65, 418–420. https://doi.org/10.2307/3280287

39. Lasonder, E., Rijpma, S. R., van Schaijk, B. C., Hoeijmakers, W. A., Kensche, P. R., Gresnigt, M. S., Sauerwein, R. W. (2016). Integrated transcriptomic and proteomic analyses of P. falciparum gametocytes: molecular insight into sex-specific processes and translational repression. Nucleic Acids Research, 44, 6087–6101. https://doi.org/10.1093/nar/gkw536

40. Li, Z., Cui, H., Guan, J., Liu, C., Yang, Z., & Yuan, J. (2021). Plasmodium transcription repressor AP2-O3 regulates sex-specific identity of gene expression in female gametocytes. EMBO Reports, 22, e51660. https://doi.org/10.15252/embr.202051660

41. Lopez-Barragan, M. J., Lemieux, J., Quinones, M., Williamson, K. C., Molina-Cruz, A., Cui, K., Su, X. Z. (2011). Directional gene expression and antisense transcripts in sexual and asexual stages of Plasmodium falciparum. BMC Genomics, 12, 587. https://doi.org/10.1186/1471-2164-12-587

42. Mair, G. R., Braks, J. A., Garver, L. S., Wiegant, J. C., Hall, N., Dirks, R. W., Waters, A. P. (2006). Regulation of sexual development of Plasmodium by translational repression. Science, 313, 667–669. https://doi.org/10.1126/science.1125129

43. Mair, G. R., Lasonder, E., Garver, L. S., Franke-Fayard, B. M., Carret, C. K., Wiegant, J. C., … Waters, A.P. (2010). Universal features of post-transcriptional gene regulation are critical for Plasmodium zygote development. PLoS Pathogens, 6, e1000767. https://doi.org/10.1371/journal.ppat.1000767

44. Miao, J., Fan, Q., Parker, D., Li, X., Li, J., & Cui, L. (2013a). Puf mediates translation repression of transmission-blocking vaccine candidates in malaria parasites. PLoS Pathogens, 9, e1003268. https://doi.org/10.1371/journal.ppat.1003268

45. Miao, J., Li, J., Fan, Q., Li, X., Li, X., & Cui, L. (2010). The Puf-family RNA-binding protein PfPuf2 regulates sexual development and sex differentiation in the malaria parasite Plasmodium falciparum. Journal of Cell Science, 123, 1039–1049. https://doi.org/10.1242/jcs.059824

46. Miao, J., Wang, Z., Liu, M., Parker, D., Li, X., Chen, X., & Cui, L. (2013b). Plasmodium falciparum: generation of pure gametocyte culture by heparin treatment. Experimental parasitology, 135, 541–545. https://doi.org/10.1016/j.exppara.2013.09.010

47. Minns, A. M., Hart, K. J., Subramanian, S., Hafenstein, S., & Lindner, S. E. (2018). Nuclear, Cytosolic, and Surface-Localized Poly(A)-Binding Proteins of Plasmodium yoelii. mSphere, 3, e00435–17. https://doi.org/10.1128/msphere.00435-17

48. Modrzynska, K., Pfander, C., Chappell, L., Yu, L., Suarez, C., Dundas, K., … Billker, O. (2017). A Knockout Screen of ApiAP2 Genes Reveals Networks of Interacting Transcriptional Regulators Controlling the Plasmodium Life Cycle. Cell Host and Microbe, 21, 11–22. https://doi.org/10.1016/j.chom.2016.12.003

49. Mora, C., McKenzie, T., Gaw, I. M., Dean, J. M., von Hammerstein, H., Knudson, T. A., … Franklin, E.C. (2022). Over half of known human pathogenic diseases can be aggravated by climate change. Nature Climate Change, 12, 869–875. https://doi.org/10.1038/s41558-022-01426-1

50. Muller, K., Silvie, O., Mollenkopf, H. J., & Matuschewski, K. (2021). Pleiotropic Roles for the Plasmodium berghei RNA Binding Protein UIS12 in Transmission and Oocyst Maturation. Frontiers in Cellular and Infection Microbiology, 11, 624945. https://doi.org/10.3389/fcimb.2021.624945

51. Munoz, E. E., Hart, K. J., Walker, M. P., Kennedy, M. F., Shipley, M. M., & Lindner, S. E. (2017). ALBA4 modulates its stage-specific interactions and specific mRNA fates during Plasmodium yoelii growth and transmission. Molecular Microbiology, 106, 266–284. https://doi.org/10.1111/mmi.13762

52. Musabyimana, J. P., Distler, U., Sassmannshausen, J., Berks, C., Manti, J., Bennink, S., … Pradel, G. (2022). Plasmodium falciparum S-Adenosylmethionine Synthetase Is Essential for Parasite Survival through a Complex Interaction Network with Cytoplasmic and Nuclear Proteins. Microorganisms, 10, 1419. https://doi.org/10.3390/microorganisms10071419

53. Ngwa, C. J., Farrukh, A., & Pradel, G. (2021). Zinc finger proteins of Plasmodium falciparum. Cellular Microbiology, 23, e13387. https://doi.org/10.1111/cmi.13387

54. Ngwa, C. J., Kiesow, M. J., Papst, O., Orchard, L. M., Filarsky, M., Rosinski, A. N., Pradel,G. (2017). Transcriptional Profiling Defines Histone Acetylation as a Regulator of Gene Expression during Human-to-Mosquito Transmission of the Malaria Parasite Plasmodium falciparum. Frontiers in Cellular and Infection Microbiology, 7, 320. https://doi.org/10.3389/fcmb.2017.00320

55. Parkyn Schneider, M., Liu, B., Glock, P., Suttie, A., McHugh, E., Andrew, D., Dixon, M. W. A. (2017). Disrupting assembly of the inner membrane complex blocks Plasmodium falciparum sexual stage development. PLoS Pathogens, 13, e1006659. https://doi.org/10.1371/journal.ppat.1006659

56. Ponzi, M., Sidén-Kiamos, I., Bertuccini, L., Currà, C., Kroeze, H., Camarda, G., … Alano, P. (2009). Egress of Plasmodium berghei gametes from their host erythrocyte is mediated by the MDV-1/PEG3 protein. Cellular Microbiology, 11, 1272–1288. https://doi.org/10.1111/j.1462-5822.2009.01331.x

57. Poran, A., Notzel, C., Aly, O., Mencia-Trinchant, N., Harris, C. T., Guzman, M. L., Kafsack, B. F. C. (2017). Single-cell RNA sequencing reveals a signature of sexual commitment in malaria parasites. Nature, 551, 95–99. https://doi.org/10.1038/nature24280

58. Prommana, P., Uthaipibull, C., Wongsombat, C., Kamchonwongpaisan, S., Yuthavong, Y., Knuepfer, E., Shaw, P. J. (2013). Inducible knockdown of Plasmodium gene expression using the glmS ribozyme. PLoS One, 8, e73783. https://doi.org/10.1371/journal.pone.0073783

59. Reddy, B. P., Shrestha, S., Hart, K. J., Liang, X., Kemirembe, K., Cui, L., & Lindner, S. E. (2015). A bioinformatic survey of RNA-binding proteins in Plasmodium. BMC Genomics, 16, 890. https://doi.org/10.1186/s12864-015-2092-1

60. Rupp, I., Sologub, L., Williamson, K. C., Scheuermayer, M., Reininger, L., Doerig, C., … Pradel, G. (2011). Malaria parasites form filamentous cell-to-cell connections during reproduction in the mosquito midgut. Cell Research, 21, 683-696. https://doi.org/10.1038/cr.2010.176

61. Russell, A. J. C., Sanderson, T., Bushell, E., Talman, A. M., Anar, B., Girling, G., … Billker,O. (2023). Regulators of male and female sexual development are critical for the transmission of a malaria parasite. Cell Host and Microbe, 31, 305–319. https://doi.org/10.1016/j.chom.2022.12.011

62. Schneweis, S., Maier, W. A., & Seitz, H. M. (1991). Haemolysis of infected erythrocytes--a trigger for formation of Plasmodium falciparum gametocytes? Parasitology Research, 77, 458–460. https://doi.org/10.1007/BF00931646

63. Shang, X., Shen, S., Tang, J., He, X., Zhao, Y., Wang, C., … Zhang, Q. (2021). A cascade of transcriptional repression determines sexual commitment and development in Plasmodium falciparum. Nucleic Acids Research, 49, 9264–9279. https://doi.org/10.1093/nar/gkab683

64. Shrestha, S., Li, X., Ning, G., Miao, J., & Cui, L. (2016). The RNA-binding protein Puf1 functions in the maintenance of gametocytes in Plasmodium falciparum. Journal of Cell Science, 129, 3144–3152. https://doi.org/10.1242/jcs.186908

65. Sielaff, M., Kuharev, J., Bohn, T., Hahlbrock, J., Bopp, T., Tenzer, S., & Distler, U. (2017). Evaluation of FASP, SP3, and iST Protocols for Proteomic Sample Preparation in the Low Microgram Range. Journal of Proteome Research, 16, 4060–4072. https://doi.org/10.1021/acs.jproteome.7b00433

66. Simon, N., Lasonder, E., Scheuermayer, M., Kuehn, A., Tews, S., Fischer, R., … Pradel, G. (2013). Malaria parasites co-opt human factor H to prevent complement-mediated lysis in the mosquito midgut. Cell Host and Microbe, 13, 29–41. https://doi.org/10.1016/j.chom.2012.11.013

67. Sinha, A., Baumgarten, S., Distiller, A., McHugh, E., Chen, P., Singh, M., … Scherf, A. (2021). Functional Characterization of the m(6)A-Dependent Translational Modulator PfYTH.2 in the Human Malaria Parasite. mBio, 12, e00661–21. https://doi.org/10.1128/mbio.00661-21

68. Sinha, A., Hughes, K. R., Modrzynska, K. K., Otto, T. D., Pfander, C., Dickens, N. J., … Billker, O. (2014). A cascade of DNA-binding proteins for sexual commitment and development in Plasmodium. Nature, 507, 253–257. https://doi.org/10.1038/nature12970

69. Smith, T. G., Lourenco, P., Carter, R., Walliker, D., & Ranford-Cartwright, L. C. (2000). Commitment to sexual differentiation in the human malaria parasite, Plasmodium falciparum. Parasitology 121 (Pt 2), 127–133. https://doi.org/10.1017/s0031182099006265

70. Sologub, L., Kuehn, A., Kern, S., Przyborski, J., Schillig, R., & Pradel, G. (2011). Malaria proteases mediate inside-out egress of gametocytes from red blood cells following parasite transmission to the mosquito. Cellular Microbiology, 13, 897–912. https://doi.org/10.1111/j.1462-5822.2011.01588.x

71. Szklarczyk, D., Gable, A. L., Lyon, D., Junge, A., Wyder, S., Huerta-Cepas, J., … Mering, C. V. (2019). STRING v11: protein-protein association networks with increased coverage, supporting functional discovery in genome-wide experimental datasets. Nucleic Acids Research, 47, D607-D613. https://doi.org/10.1093/nar/gky1131

72. Tarique, M., Ahmad, M., Ansari, A., & Tuteja, R. (2013). Plasmodium falciparum DOZI, an RNA helicase interacts with eIF4E. Gene, 522, 46–59. https://doi.org/10.1016/j.gene.2013.03.063

73. Usui, M., Prajapati, S. K., Ayanful-Torgby, R., Acquah, F. K., Cudjoe, E., Kakaney, C., … Williamson, K. C. (2019). Plasmodium falciparum sexual differentiation in malaria patients is associated with host factors and GDV1-dependent genes. Nature Communications, 10, 2140. https://doi.org/10.1038/s41467-019-10172-6

74. van Biljon, R., van Wyk, R., Painter, H. J., Orchard, L., Reader, J., Niemand, J., Birkholtz, L. M. (2019). Hierarchical transcriptional control regulates Plasmodium falciparum sexual differentiation. BMC Genomics, 20, 920. https://doi.org/10.1186/s12864-019-6322-9

75. Varadi, M., Anyango, S., Deshpande, M., Nair, S., Natassia, C., Yordanova, G., Velanker, S. (2022). AlphaFold Protein Structure Database: massively expanding the structural coverage of protein-sequence space with high-accuracy models. Nucleic Acids Research, 50, D439–D444. https://doi.org/10.1093/nar/gkab1061

76. Vembar, S. S., Macpherson, C. R., Sismeiro, O., Coppee, J. Y., & Scherf, A. (2015). The PfAlba1 RNA-binding protein is an important regulator of translational timing in Plasmodium falciparum blood stages. Genome Biology, 16, 212. https://doi.org/10.1186/s13059-015-0771-5

77. Vizcaino, J. A., Cote, R. G., Csordas, A., Dianes, J. A., Fabregat, A., Foster, J. M., … Hermjakob, H. (2013). The PRoteomics IDEntifications (PRIDE) database and associated tools: status in 2013. Nucleic Acids Research, 41, D1063–D1069. https://doi.org/10.1093/nar/gks1262

78. Weißbach, T., Golzmann, A., Bennink, S., Pradel, G., & Ngwa, C. J. (2017). Transcript and protein expression analysis of proteases in the blood stages of Plasmodium falciparum. Experimental Parasitology, 180, 33–44. https://doi.org/10.1016/j.exppara.2017.03.006

79. Wirth, C. C., Glushakova, S., Scheuermayer, M., Repnik, U., Garg, S., Schaack, D., … Pradel, G. (2014). Perforin-like protein PPLP2 permeabilizes the red blood cell membrane during egress of Plasmodium falciparum gametocytes. Cellular Microbiology, 16, 709–733. https://doi.org/10.1111/cmi.12288

80. WHO World Malaria Report (2022). https://www.who.int/teams/global-malaria-programme/reports/world-malaria-report-2022

81. Xu, Y., Qiao, D., Wen, Y., Bi, Y., Chen, Y., Huang, Z., Cui, L., Guo, J., Cao, Y. (2021). PfAP2-G2 is associated to production and maturation of gametocytes in Plasmodium falciparum via regulating the expression of PfMDV-1. Frontiers in Microbiology, 11, 631444. https://doi.org/10.3389/fmicb.2020.631444

82. Yuda, M., Iwanaga, S., Kaneko, I., & Kato, T. (2015). Global transcriptional repression: An initial and essential step for Plasmodium sexual development. Proceedings of the National Academy of Sciences USA, 112, 12824–12829. https://doi.org/10.1073/pnas.1504389112

83. Yuda, M., Kaneko, I., Iwanaga, S., Murata, Y., & Kato, T. (2020). Female-specific gene regulation in malaria parasites by an AP2-family transcription factor. Molecular Microbiology, 113, 40–51. https://doi.org/10.1111/mmi.14334

84. Zhang, C., Li, Z., Cui, H., Jiang, Y., Yang, Z., Wang, X., … Yuan, J. (2017). Systematic CRISPR-Cas9-Mediated Modifications of Plasmodium yoelii ApiAP2 Genes Reveal Functional Insights into Parasite Development. mBio 8, e01986–17. https://doi.org/10.1128/mbio.01986-17

