## Supplemental Figures for "The *Plasmodium falciparum* CCCH zinc finger protein MD3 regulates male gametocytogenesis through its interaction with RNA-binding proteins"

*Original Research Article*

Supplemental Figures S1-S14

Figure S1

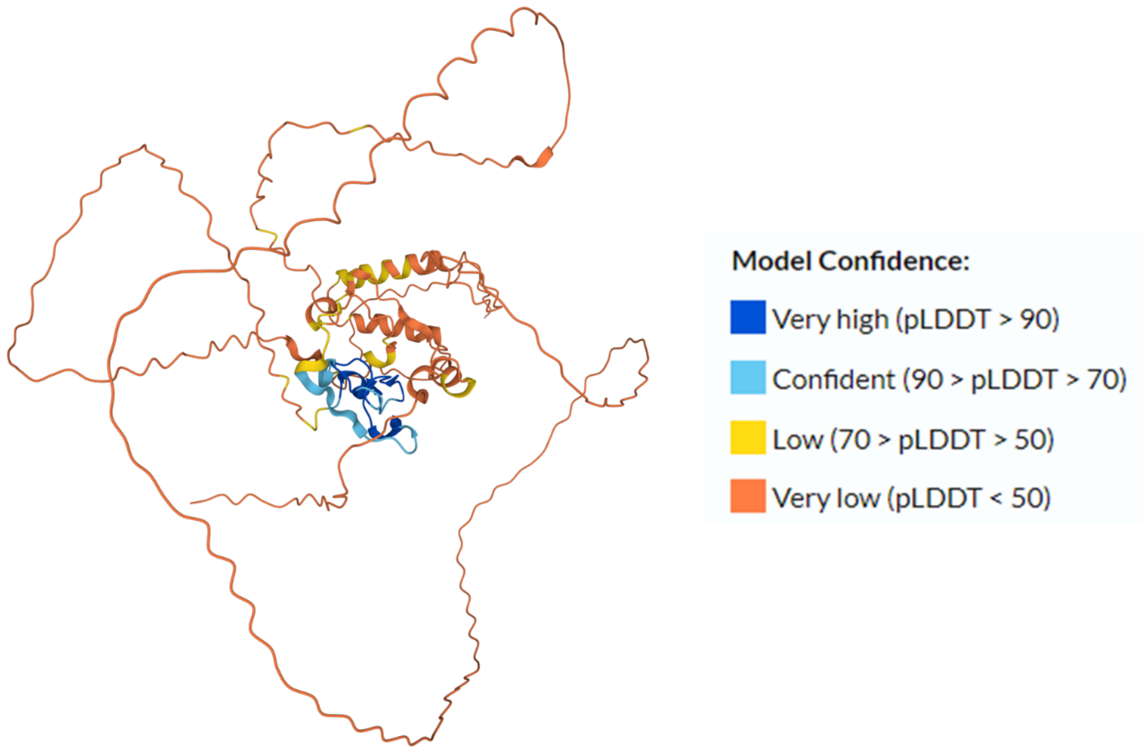


**Figure S1: Predicted 3D structure of MD3.** The 3D structure of the C3H1-ZFP MD3 was predicted, using the AlphaFold programme. Model confidence is shown by colour codes.

Figure S2

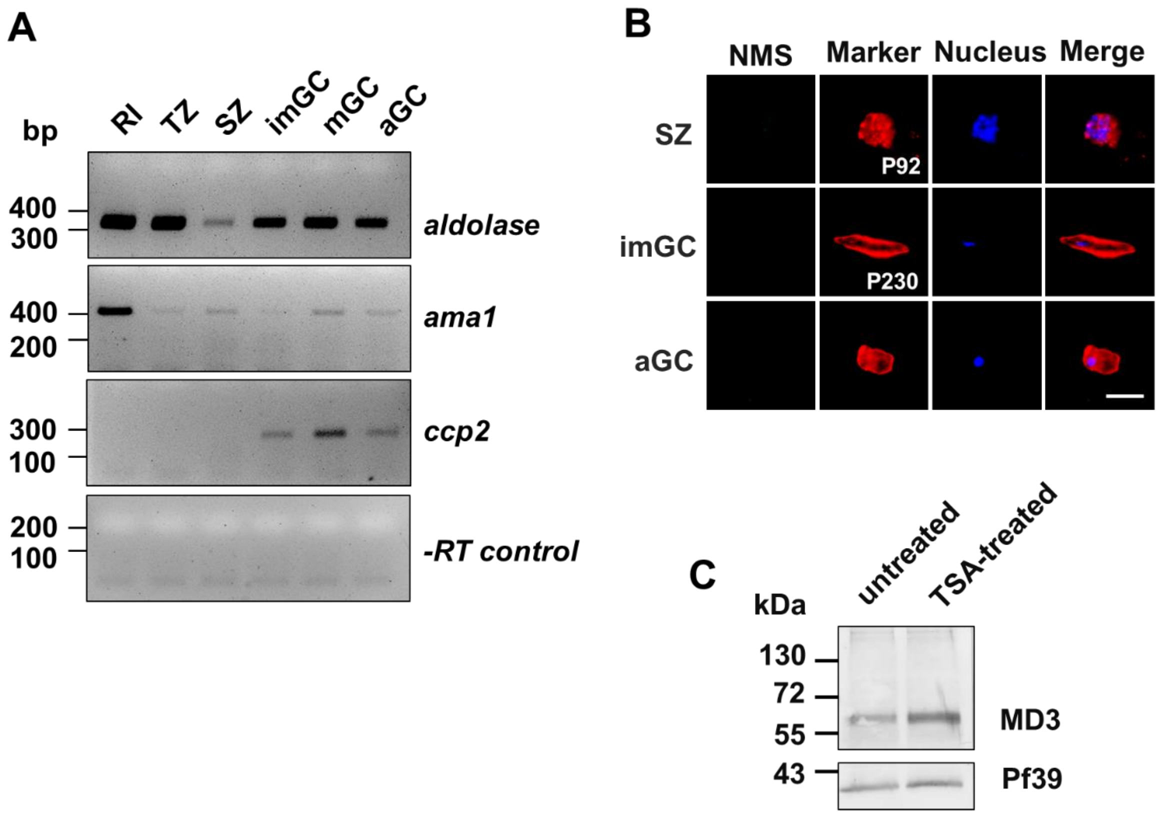


**Figure S2: MD3 transcript and protein expression controls. (A)** RT-PCR control. Diagnostic RT-PCR was used to amplify transcript of control genes from cDNA generated from total RNA of rings (RI), trophozoites (TR), schizonts (SZ), as well as immature gametocytes (imGC; stages II-IV), mature gametocytes (mGC; stage V) and activated gametocytes (aGC; 30' post-activation) of WT NF54. Transcript amplification of *ama1* (407 bp) and *ccp2* (286 bp) was used to validate the specificity of the asexual blood stage and gametocytes samples, respectively. Samples lacking reverse transcriptase (-RT) were used to investigate potential contamination with gDNA. Transcript amplification of *aldolase* (378 bp) was used as a positive control. **(B)** IFA controls. Schizonts, immature gametocytes and activated gametocytes were subjected to IFA and labelled with serum from non-immunized mice (NMS; green). SZ was counterlabelled with rabbit anti-P92 antibody and gametocytes with rabbit anti-P230 antisera (red); nuclei were highlighted by Hoechst 33342 nuclear stain (blue). Bar, 5 µm. **(C)** MD3 levels following TSA treatment. Immature gametocytes were treated with 0.26 µM TSA or 0.5% (v/v) ethanol for control for 24 h. Lysates were subjected to WB, using mouse anti-MD3 antisera to detect MD3 with the expected molecular weight of ~57 kDa. Immunoblotting with rabbit antisera against *Pf*39 (~39 kDa) served as loading and standardization control. Results (A-C) are representative of two to three independent experiments.

Figure S3

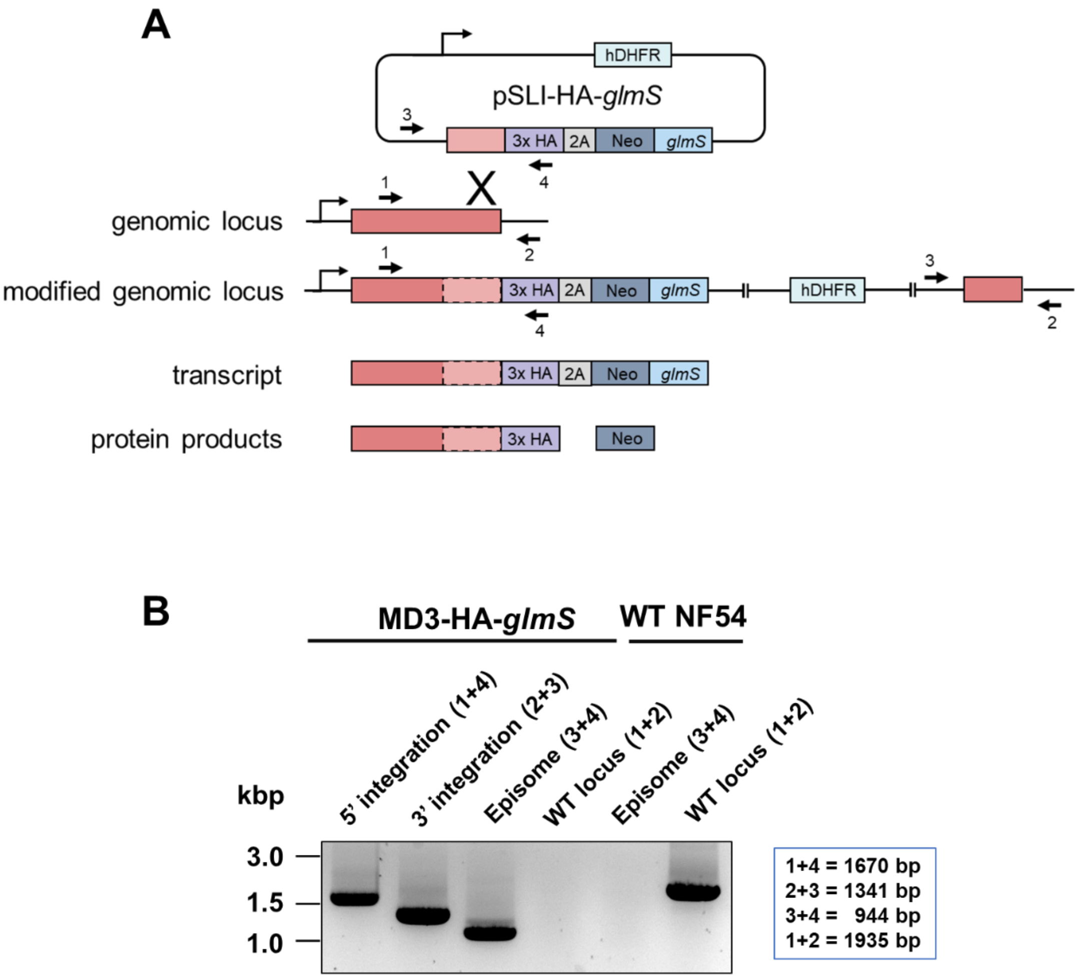


**Figure S3: Generation of the MD3-HA-*glmS* parasite line. (A)** Schematic depicting the single-crossover homologous recombination strategy for the generation of the pSLI-HA-*glmS*-based MD3-HA-*glmS* line. The coding region of the gene of interest was fused at the 3'-end to a HA-encoding sequence followed by the 2A-skip peptide sequence and the Neo and *glmS*-ribozyme sequences. The numbered arrows indicate the positions of primers used to confirm vector integration. *glmS*, glucosamine-6-phosphate-activated ribozyme; HA, hemagglutinin; hDHFR, human dihydrofolate reductase-encoding gene conferring resistance to WR99210; Neo, gene conferring resistance to neomycin. **(B)** Confirmation of vector integration into the *md3* gene locus. Diagnostic PCR demonstrates successful 5′ (primers 1 and 4) and 3′ (primers

3 and 2) integration in line MD3-HA-*glmS*. As a control, WT NF54 gDNA was used, demonstrating the original gene locus (primers 1 and 2). Episomal DNA was further detected (primers 3 and 4). Band sizes are indicated. Primer sequences are provided in Table S5.

Figure S4

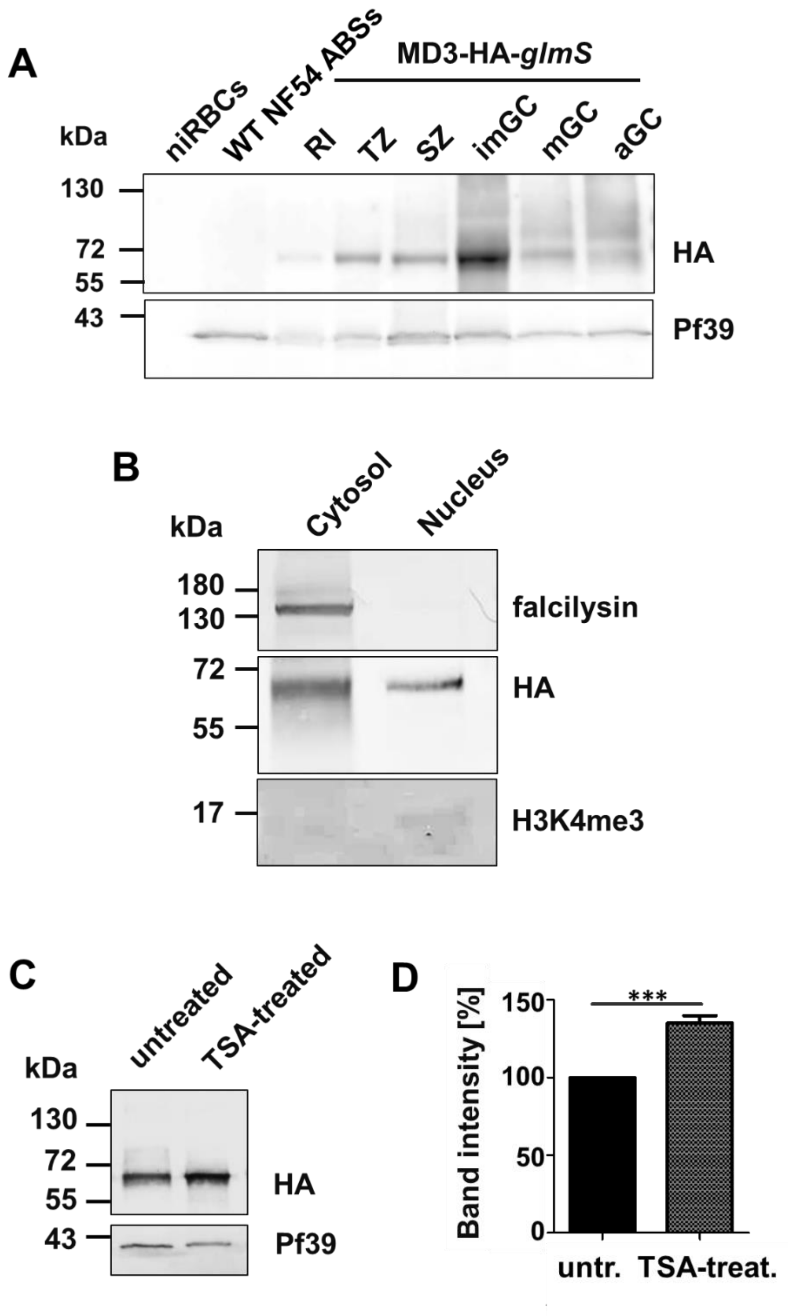


**Figure S4: MD3-HA expression and epigenetic regulation in blood-stage parasites. (A)** Expression of MD3-HA in blood stage parasites. Lysates from rings (RI), trophozoites (TZ), schizonts (SZ), immature gametocytes (imGC), mature gametocytes (mGC), and gametocytes (aGC; 30 min post-activation) of line MD3-HA-*glmS* were immunoblotted with rat anti-HA antibody to detect MD3-HA running at the expected molecular weight of ~60 kDa. Lysates of niRBCs and mixed asexual blood stages (ABS) of WT NF54 were used as negative control. Immunoblotting with rabbit anti-*Pf*39 antisera (~39 kDa) served as loading control. **(B)** Subcellular localization of MD3-HA. Cytosolic and nuclear protein fractions were extracted from immature gametocytes of line MD3-HA-*glmS* and immunoblotted with rat anti-HA antibody to detect MD3-HA (~60 kDa). Mouse antisera against the cytosolic protease falcilysin (~138 kDa) and rabbit antisera against the histone H3 mark H3K4me3 (~15 kDa) were used to confirm the fraction purity. **(C)** MD3-HA levels following TSA treatment. Immature gametocytes of line MD3-HA-*glmS* were treated with 0.26 µM TSA or 0.5% (v/v) ethanol for control for 24 h. Lysates were subjected to WB, using mouse anti-MD3 antisera to detect MD3 (~60 kDa). Immunoblotting with rabbit antisera against *Pf*39 (~39 kDa) served as loading and standardization control. **(D)** Quantification of MD3-HA levels following TSA-treatment. Lysates of TSA-treated and untreated immature gametocytes of line MD3-HA-*glmS* were prepared and subjected to WB, using anti-MD3 antisera as described in (C). MD3 protein levels were evaluated by measuring the band intensities in three independent WB experiments using Image J; the values were normalized to the respective *Pf*39 protein band, used as loading control. Results are shown as mean ± SD (untreated set to 100%). ***, $p < 0.001$ (Student's t-test). Results (A-C) are representative of two to three independent experiments.

Figure S5

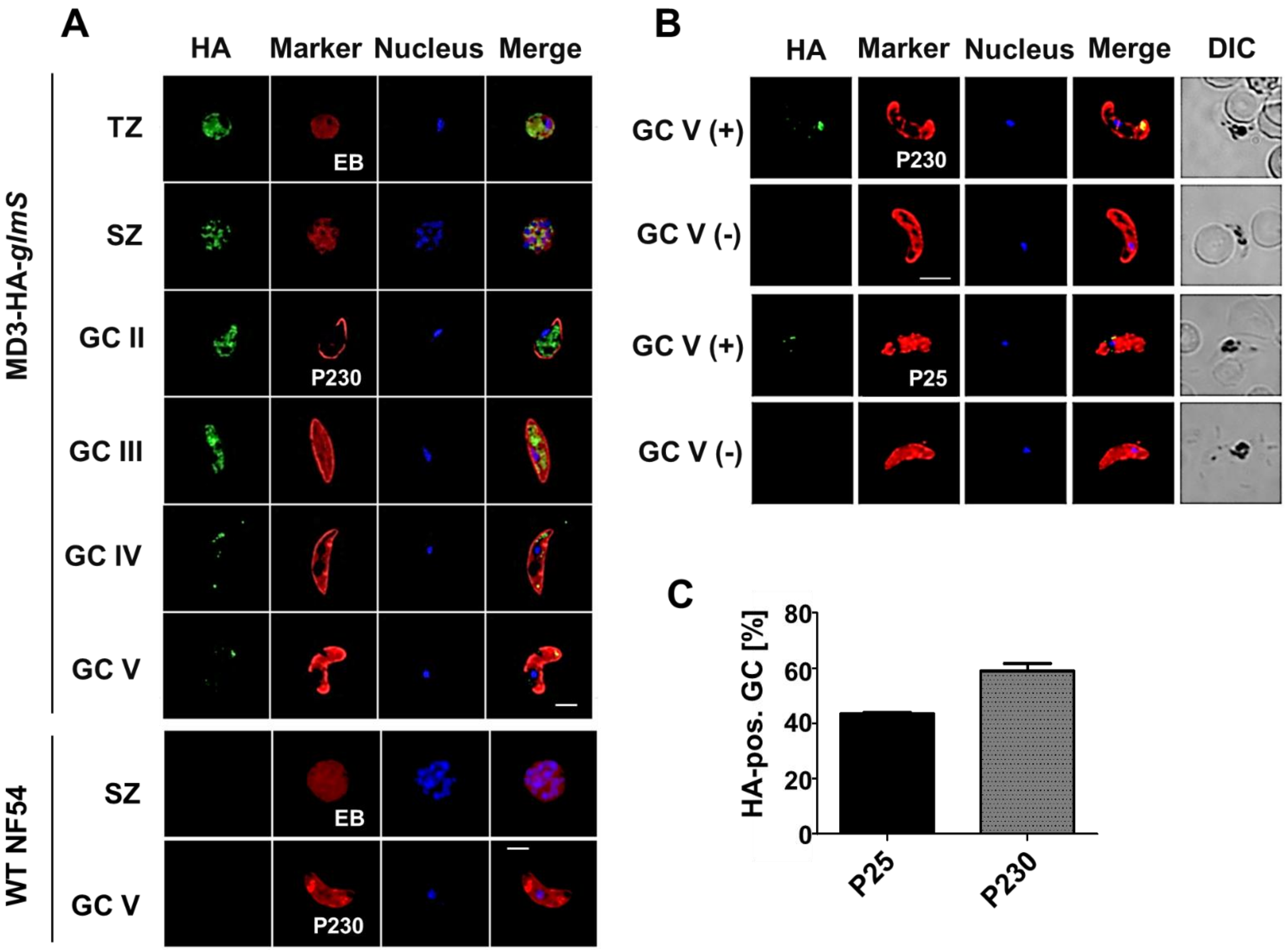


**Figure S5: Localization and sex specificity of MD3-HA in blood stage parasites. (A)** Localization of MD3-HA in blood stage parasites. Methanol-fixed rings (RI), trophozoites (TZ), schizonts (SZ), and gametocyte (GC) stages II-V of line MD3-HA-*glmS* were immunolabeled with rat anti-HA antibody (green). Asexual blood stages were stained with Evans Blue, while gametocytes were counterlabelled with rabbit anti-P230 antisera (red); nuclei were highlighted by Hoechst 33342 nuclear stain (blue). Bar, 5 µm. WT NF54 was used as a negative control. **(B)** Sex-specific expression of MD3-HA. Methanol-fixed mature gametocytes of line MD3-HA-*glmS* were immunolabeled with rat anti-HA (green) and counterlabelled (red) with either rabbit antisera against P230 (expressed in male and female gametocytes) or P25 (expressed in female gametocytes). Nuclei were stained by Hoechst 33342 nuclear stain (blue). Bar, 5 µm. **(C)** Sex specificity of MD3-HA. Mature gametocytes of line MD3-HA-*glmS* were immunolabeled as described in (B). For each marker, 100 labelled gametocytes were counted and the percentage of MD3-HA-positive gametocytes was calculated. The experiment was performed in triplicate (mean ± SD). Results (A, B) are representative of three independent experiments.

Figure S6

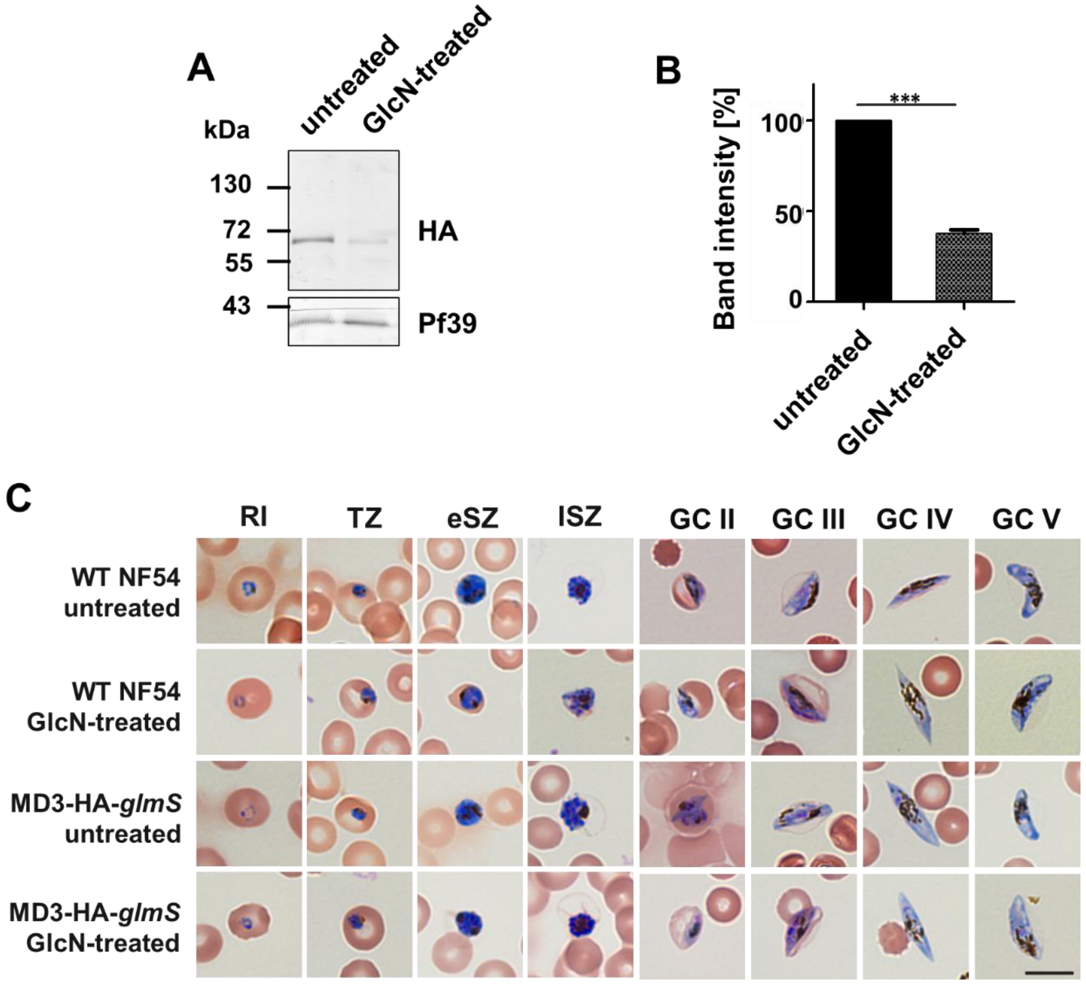


**Figure S6: Verification of MD3-HA knockdown and effect of MD3-HA knockdown on blood stage morphology. (A)** Confirmation of MD3-HA knockdown. Lysates of mixed asexual blood stages of line MD3-HA-*glmS* treated or not with 5 mM GlcN for 72 h were subjected to WB analysis using rat anti-HA antibody to detect MD3-HA (~60 kDa). Equal loading was confirmed by immunoblotting with rabbit antisera against *Pf*39 (~39 kDa). **(B)** Quantification of MD3-HA levels following knockdown. MD3-HA specific protein band intensities of three independent WB experiments as described in (A) were quantified using Image J and normalized to the respective *Pf*39 protein band intensities (untreated set to 100%). Results are shown as mean ± SD. ***, $p < 0.001$ (Student's t-test). **(C)** Blood stage morphology following MD3-HA knockdown. Rings (RI), trophozoites (TZ), early schizonts (eSZ), late schizonts (lSZ) and gametocyte (GC) stages II-V of line MD3-HA-*glmS* and WT NF54 treated or not with 5 mM GlcN for 72 h were Giemsa stained to compare their morphologies. Bar, 5 μm.

Figure S7

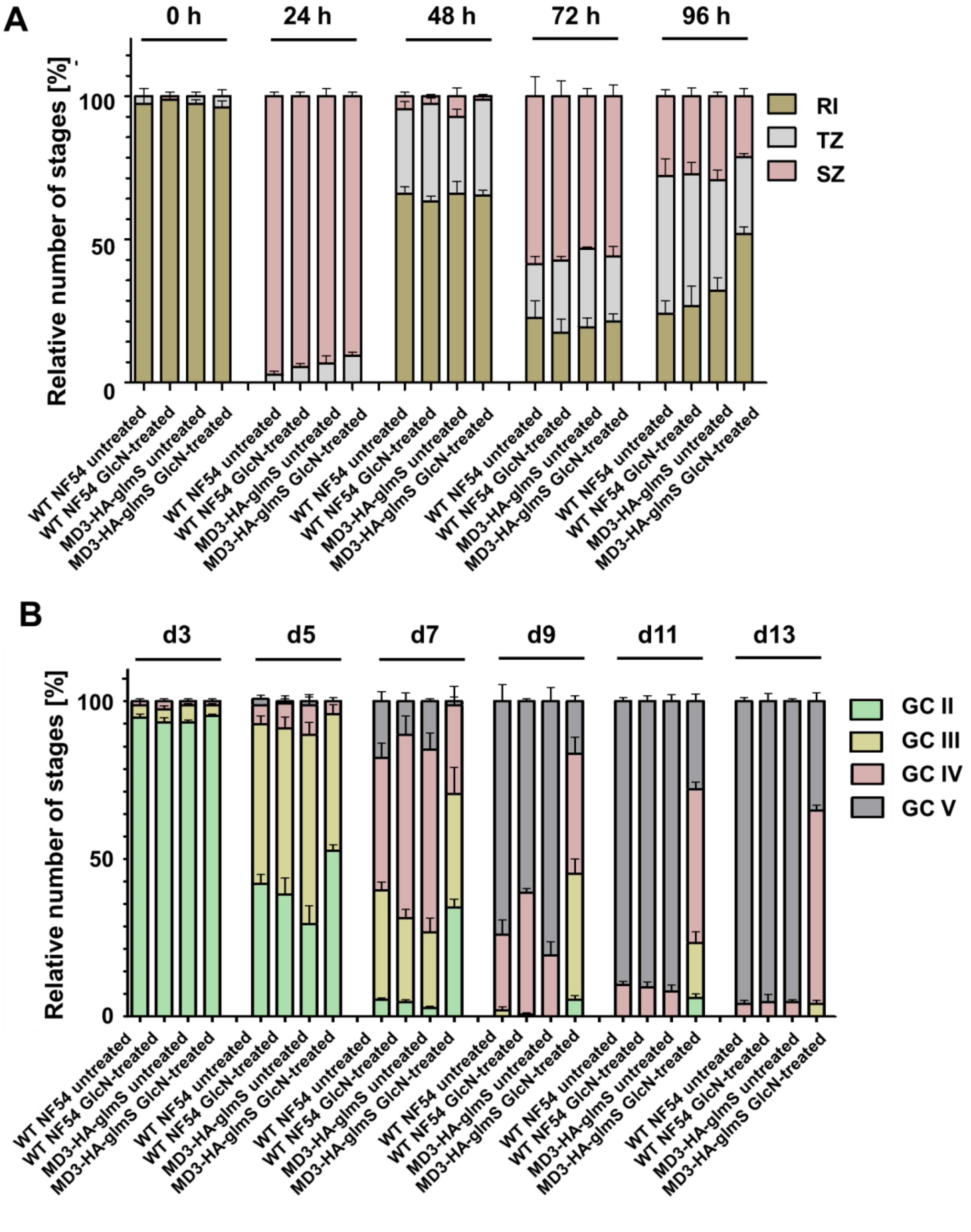


**Figure S7: Stage development in blood stage parasites following MD3-HA knockdown.** **(A)** Asexual blood stage development following MD3-HA knockdown. Synchronized ring stage cultures of WT NF54 and line MD3-HA-*glmS* with a starting parasitaemia of 0.25% were maintained in cell culture medium supplemented or not with 5 mM GlcN for transcript knockdown over a time-period of 0 - 96 h. Rings (RI), trophozoites (TR), and schizonts (SZ) were determined in a total number of 50 infected RBCs every 24 h via Giemsa-stained blood smears. The experiment was performed in triplicate. **(B)** Gametocyte maturation following MD3-HA knockdown. Gametocyte production was induced in synchronized ring stage cultures of WT NF54 and line MD3-HA-*glmS* at a final parasitaemia of 7.5% by addition of lysed RBCs. Subsequently, the cultures were maintained in cell culture medium supplemented or not with 5 mM GlcN for transcript knockdown for 13 days. Relative numbers of gametocyte stages II-V were followed via Giemsa smears in 50 iRBCs every 48 hours. The experiment was performed in triplicate.

Figure S8

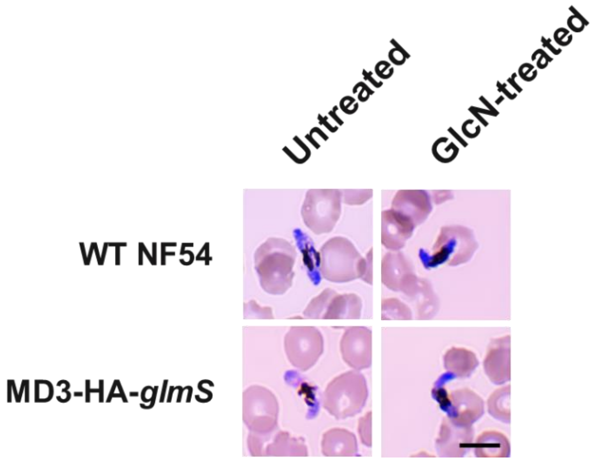


**Figure S8: Giemsa-stained images of mature gametocytes used for the exflagellation assays.** Gametocyte production was induced in synchronized ring stage cultures of WT NF54 and line MD3-HA-*glmS* at a final parasitaemia of 7.5% by addition of lysed RBCs. Following treatment with 20 U/ml of heparin for 2 days, cultures were maintained in normal cell culture medium. On day 7-10 post-induction, the gametocytes were maintained in cell culture medium supplemented or not with 5 mM GlcN for transcript knockdown. On day 13, Giemsa smears were performed to determine the numbers of mature gametocytes in the cultures. Bar, 5 µm.

Figure S9

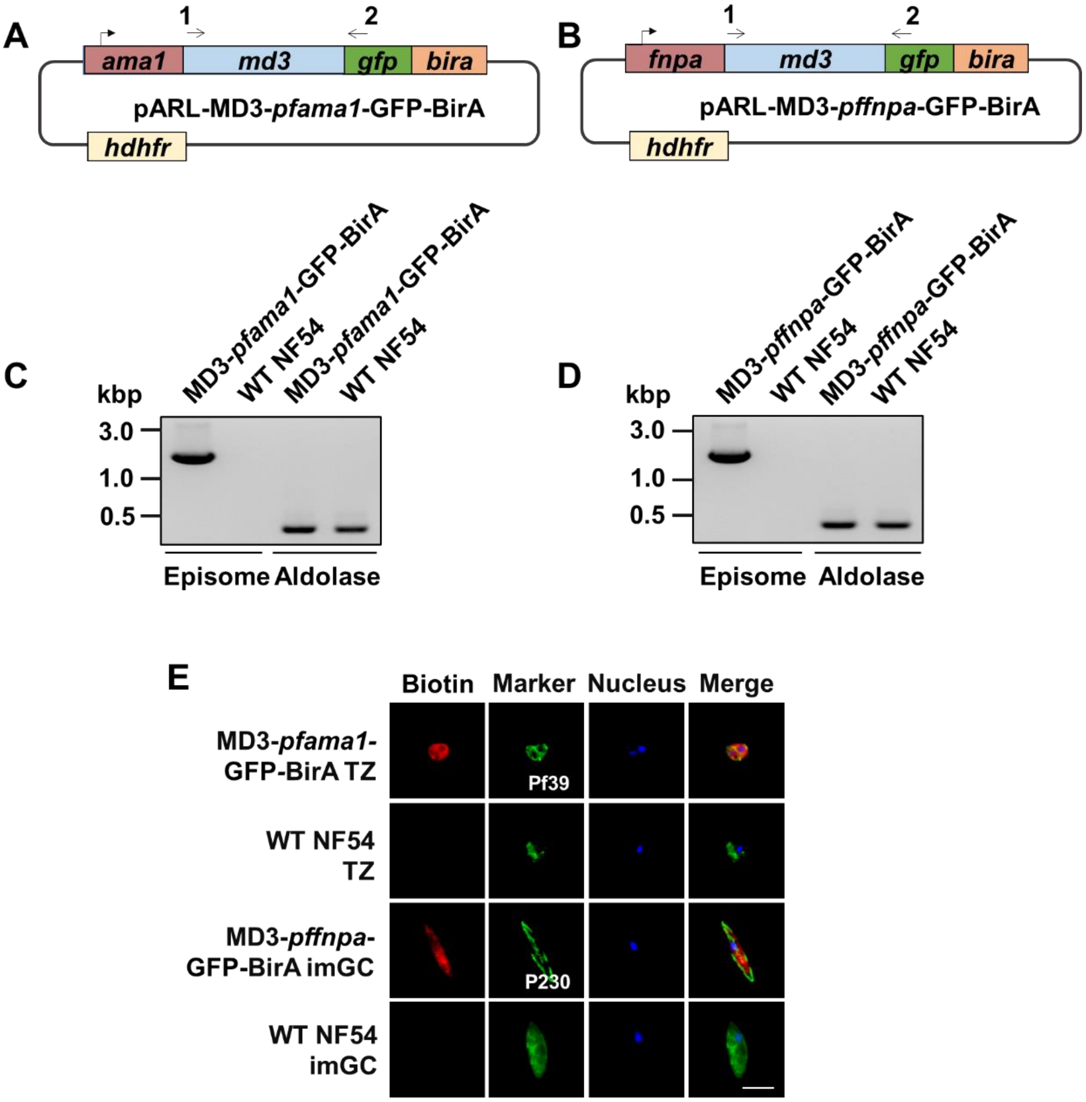


**Figure S9: Generation of the MD3-GFP-BirA parasite lines. (A, B)** Schematics depicting the vectors pARL-MD3-*pfama1*-BirA and pARL-MD3-*pffnpa*-GFP-BirA, comprising the *pfama1* and *pffnpa* promoters, respectively. BirA, *E. coli* biotin ligase; GFP, green fluorescent protein; *hdhfr*, human dihydrofolate reductase encoding gene referring resistance to WR99210. **(C, D)** Verification of vector uptake in the MD3-GFP-BirA lines. Diagnostic PCR was employed to detect the presence of vectors pARL-MD3-*pfama1*-BirA and pARL-MD3-*pffnpa*-GFP-BirA (primers 1 and 2; 1687 bp) in lines MD3-*pfama1*-BirA and MD3-*pffnpa*-GFP-BirA, but not the WT NF54. Transcript amplification of plasmodial *aldolase* (378 bp) was used as a control. (E) Localization of biotinylated proteins in the MD3-GFP-BirA lines. Methanol-fixed trophozoites (TZ) of MD3-*pfama1*-GFP-BirA and immature gametocytes (imGC) of MD3-*pffnpa*-GFP-BirA, which had been treated with 50 μM biotin for 24 h, were immunolabeled with streptavidin-594 (red). Asexual blood stages were counterlabelled with rabbit anti-*Pf*39 antisera and gametocytes with rabbit anti-P230 antisera (green); nuclei were highlighted by Hoechst 33342 nuclear stain (blue). WT NF54 parasites were used for negative control. Bar, 5 μm.

Figure S10

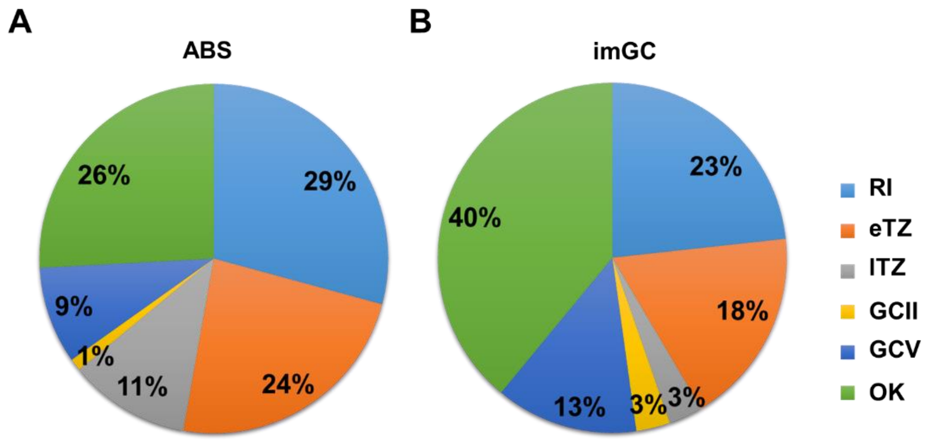


**Figure S10: Pie charts depicting the peak transcript expression of the MD3 interactors. (A, B)** The 41 MD3 interactors in the asexual blood stages (ABS; A) and the 98 MD3 interactors in immature gametocytes (imGC; B) were grouped (percentage of total numbers) by stages of peak expression (see PlasmoDB database). RI, ring stage; eTZ, early trophozoite; lTZ, late trophozoite; SZ, schizont; GC II, gametocyte stage II; GC V, gametocyte stage V; OK, ookinete.

Figure S11

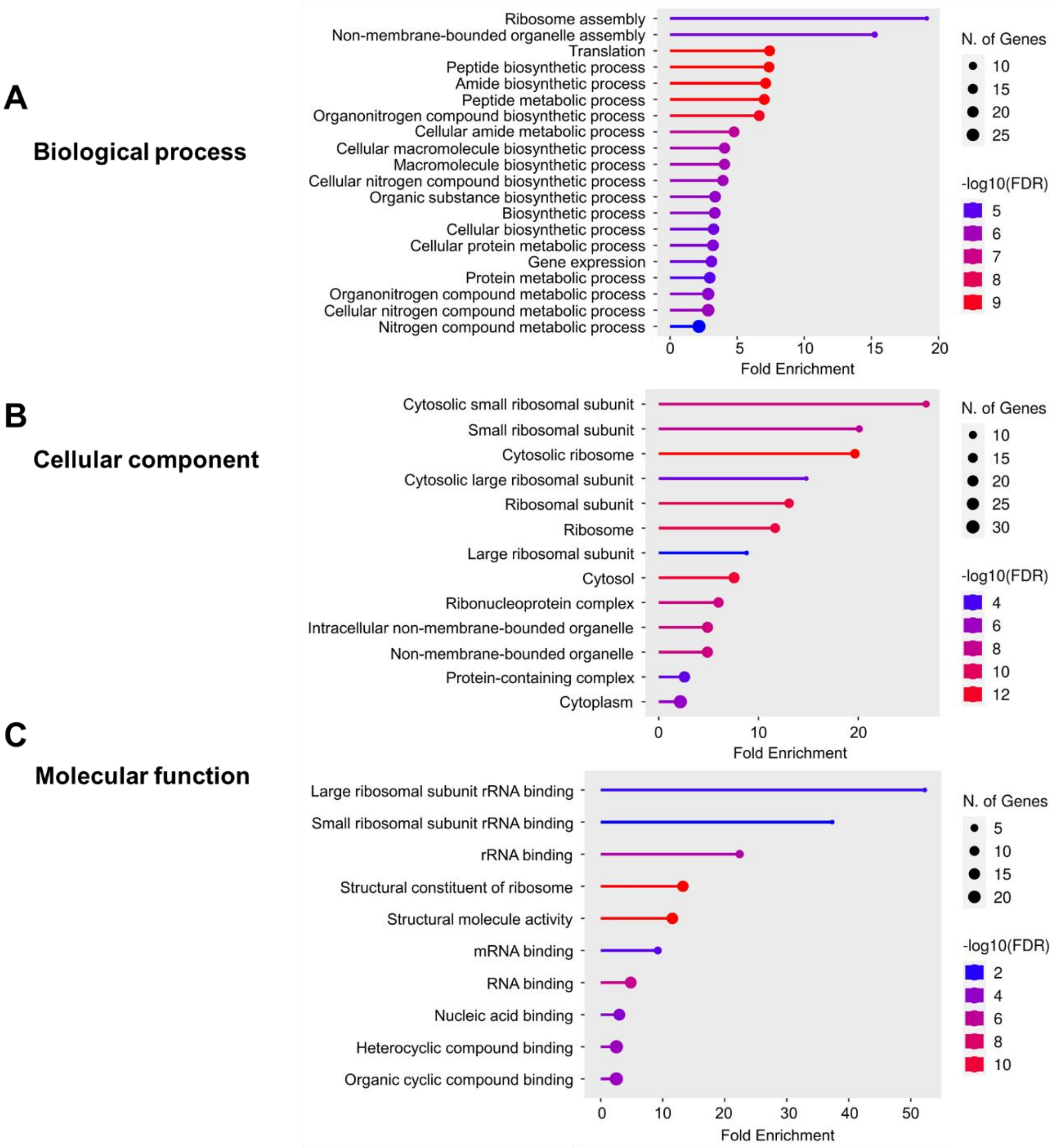


**Figure S11: Functional prediction analysis of the MD3 interactors in the asexual blood stages.** A GO enrichment analysis of the 41 putative MD3 interactors in the asexual blood stages was performed using the ShinyGO 0.77 programme with $p < 0.05$ to display the enriched GO terms based on biological process **(A)**, cellular component **(B)** and molecular function **(C)**.

Figure S12

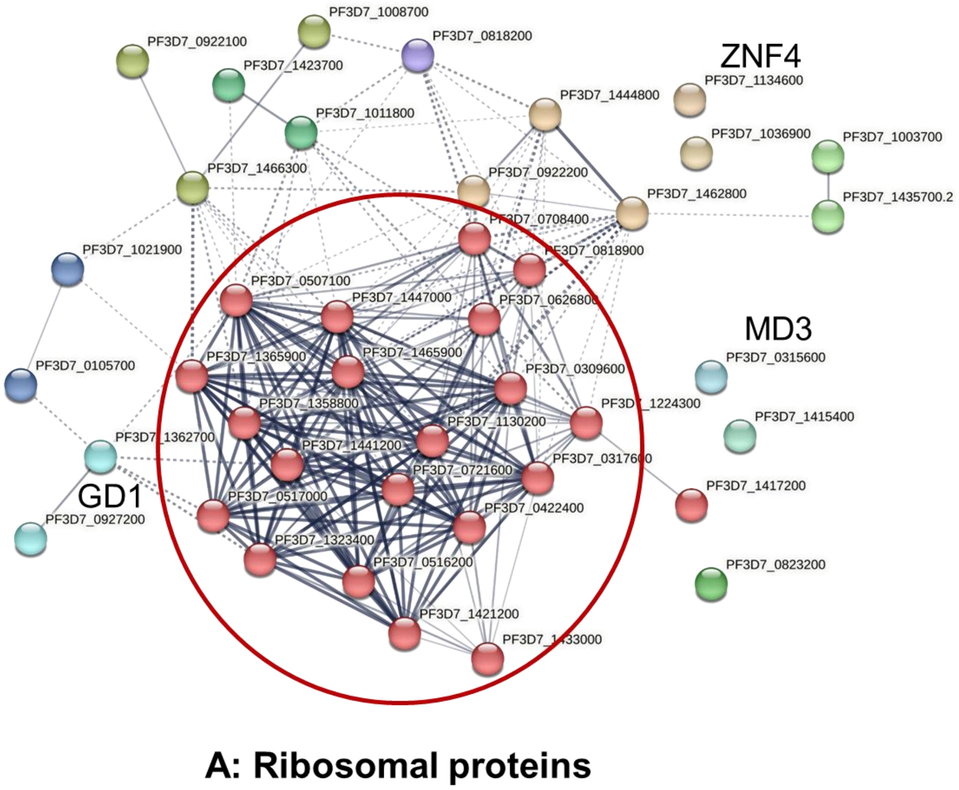


**Figure S12: Network analysis of the MD3 interactors in the asexual blood stages.** A protein-protein network analysis of the 41 putative MD3 interactors in the asexual blood stages was performed, using the STRING database, selecting a medium interaction confidence of 0.4; line thickness specifies the strength of data support between the proteins. Clustering of the interactors was employed by the Markov Clustering (MCL) algorithm with an inflation parameter of 3; dotted lines represent the connection among the hits of different clusters. On the basis of physical interactions among the query proteins, one main cluster was identified, i.e. A) ribosomal proteins. Detailed information on the cluster and its components is provided in Table S4.

Figure S13

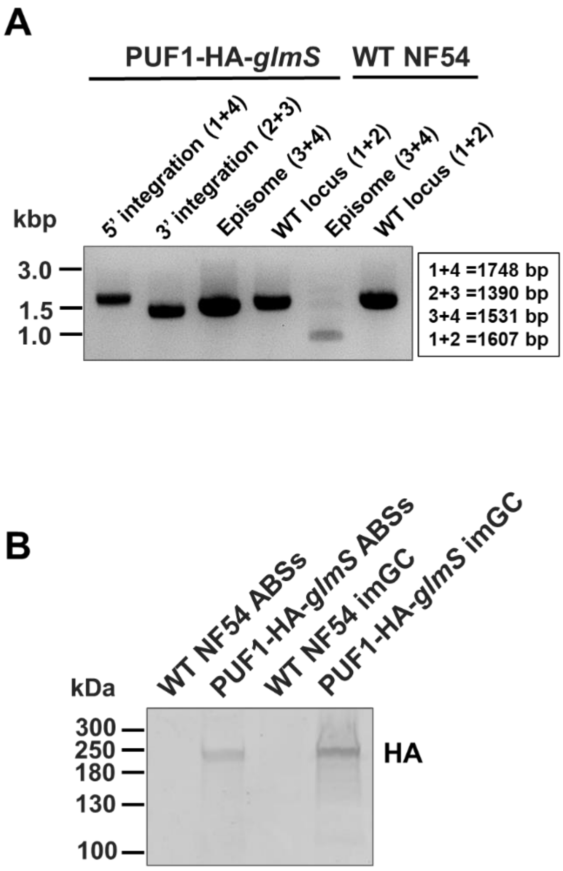


**Figure S13: Generation and verification of the PUF1-HA-*glmS* parasite line. (A)** Confirmation of vector integration into the *puf1* gene locus. Diagnostic PCR demonstrates successful 5′ (primers 1 and 4) and 3′ (primers 3 and 2) integration in line PUF1-HA-*glmS*. As a control, WT NF54 gDNA was used, demonstrating the original gene locus (primers 1 and 2). Episomal DNA was further detected (primers 3 and 4). Band sizes are indicated. Primer sequences are provided in Table S5. **(B)** Expression of PUF1-HA in the blood stages of line PUF1-HA-*glmS*. Lysates from asexual blood stages (ABS) and immature gametocytes (imGC), of line PUF1-HA-*glmS* were immunoblotted with rat anti-HA antibody to detect PUF1-HA running at the expected molecular weight of ~227 kDa. WT NF54 was used as a negative control. Results (A, B) are representative of two to three independent experiments.

Figure S14

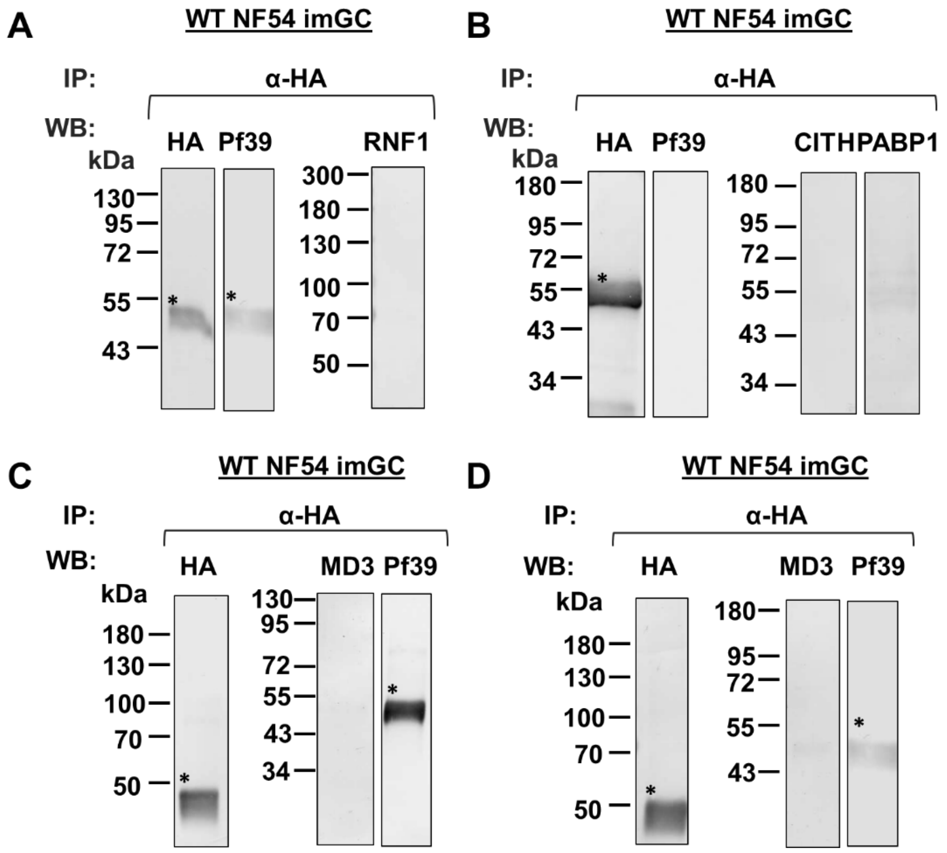


**Figure S14: Co-immunoprecipitation controls of the interactions of MD3 with RNA-binding proteins**: Lysates of immature gametocytes (imGC) of WT NF54 were subjected to co-immunoprecipitation assays. Protein complexes were immunoprecipitated using polyclonal rabbit or rat anti-HA antibodies, followed by immunoblotting using rabbit or rat anti-HA antibodies, mouse anti-RNF1 antisera, rabbit anti-CITH antibody, rabbit anti-PABP1 antibody or mouse anti-MD3 antisera to detect the precipitated proteins (expected molecular weights: MD3-HA, ~60 kDa; ZNF4-HA, ~207 kDa, PUF1-HA, ~227 kDa; RNF1, ~138 kDa, MD3, ~57 kDa, CITH, ~39 kDa, PABP1, ~97 kDa). Immunoblotting with rabbit anti-*Pf*39 served as a negative control (~39 kDa). **(A, B)**, controls for line MD3-HA-*glmS*; **(C)**, control for line ZNF4-HA-*glmS*; **(D)**, control for line PUF1-HA-*glmS*. Asterisks signify bands corresponding to the heavy chains of the precipitation antibody. Results are representative of two to three independent experiments.
