## Supplemental Table S5 for "The *Plasmodium falciparum* CCCH zinc finger protein MD3 regulates male gametocytogenesis through its interaction with RNA-binding proteins"

**Table S5. List of primers.**

| Function | Name | Sequence 5'-3' |
|---|---|---|
| **Oligonucleotides for diagnostic RT PCR** | | |
| RT Primers | MD3-RT FP | GGGTTCACGTTGCAGATTTT |
| | MD3-RT RP | TTGCGTTGTCTTTTGTTTGC |
| | Aldolase-RT FP | TAGATGGATTAGCAGAAAGATGC |
| | Aldolase-RT RP | TCAAAATCACGCACATCCTG |
| | Ama1-RT FP | CGGTAGCTACGGGAAATCAA |
| | Ama1-RT RP | AGGGCAAACTTTTTCCCAGT |
| | CCp2-RT FP | AGTTGTTGATGGGCTTTTGG |
| | CCp2-RT RP | ATTCGGTGCCATTAGGGTTA |
| **Oligonucleotides for the generation of MD3 and RNF1 recombinant proteins** | | |
| Cloning primers | MD3 NotI FP | agatctgcggccgcTGGGTCGACTTACCCGAAACAA |
| | MD3 PstI RP | agatctctgcagCATTTGATTTCCCCTTGGACAG |
| | RNF1 NotI FP | agatctgcggccgcATCGCACAGAAGCCTAAGGA |
| | RNF1 PstI RP | agatctctgcagTCCCATATTTCGCATTAGCC |
| **Oligonucleotides for the generation and verification of MD3-HA-glmS mutant** | | |
| Cloning Primers | MD3-HA-glmS NotI FP | tcctccgcggccgcCGCAAGGGAGGGGAAAATAAA |
| | MD3-HA-glmS XmaI RP | ctttactcccgggGTGTCTCCTCATGGGTGCCATGT |
| Integration Primers | pSLI-MD3-HA-glmS FP | GAGGAAACTTTTTATTCACATCTGAA |
| | pSLI-MD3-HA-glmS RP | CCACATACGCTTAATTATAATATGATG |
| | pARL-HA-glmS FP | GCTTTACACTTTATGCTTCCGGCTC |
| | pSLI-HA-glmS RP | TGTCTGTTGTGCCCAGTCAT |

| | | |
|---|---|---|
| **Oligonucleotides for the generation and verification of MD3-*pfama1*/*pffnpa*-GFP-BirA mutants** | | |
| Cloning primers | MD3-GFP-BirA KpnI FP | atgcatggtaccATGAATTCAAATCATAATATGTTAA |
| | MD3-GFP-BirA AvrII RP | atgcatcctaggGTGTCTCCTCATGGGTGCCA |
| Diagnostic PCR Primers | MD3-GFP-BirA KpnI FP | atgcatggtaccATGAATTCAAATCATAATATGTTAA |
| | pGREP GFP RP | CAAGTGTTGGCCATGGAA |
| | Aldolase-RT FP | TAGATGGATTAGCAGAAAGATGC |
| | Aldolase-RT RP | TCAAAATCACGCACATCCTG |
| **Oligonucleotides for the generation and verification of PUF1-HA-glmS mutant** | | |
| Cloning Primers | PUF1-HA-glmS NotI FP | tcctccgcggccgcTTGGTCAAAACTTGCATAGGG |
| | PUF1-HA-glmS XmaI RP | ctttactcccgggTATTAATTTGGACTCATTACTATATTTCTTC |
| Integration Primers | pSLI-PUF1-glmS FP | TCCCCACAAAATATTGCACAT |
| | pSLI-PUF1-glmS RP | TTTTGTTTGTCTAATCATTTGCC |
| | pARL-HA-glmS FP | GCTTTACACTTTATGCTTCCGGCTC |
| | pSLI-HA-glmS RP | TGTCTGTTGTGCCCAGTCAT |

FP, forward primer; RP, reverse primer; RT, reverse transcriptase; restriction sites indicated in red colour
